# Spatial and lineage dependent processes underpin floristic assembly in the megadiverse Eastern South American mountains

**DOI:** 10.1101/2022.05.26.493493

**Authors:** Yago Barros-Souza, Leonardo M. Borges

## Abstract

**Aim:** The astonishing diversity of ancient mountains was likely shaped by multiple evolutionary processes. However, there is an ongoing debate on what were the main processes driving the assembly of *campos rupestres*, the mega-diverse flora of Eastern South American mountains. Although the ancient nature of these mountains suggests their flora should be composed by relatively older lineages, they harbour a number of recently diverged clades. To better understand the evolution of ancient mountains’ floras, we tested if the *campos rupestres* are mainly composed by relatively old or recent communities and if angiosperm diversity is geographically structured using analyses of diversity and endemism.

**Location:** Eastern South America.

**Time period:** Oligocene/Miocene to the present.

**Major taxa studied:** Flowering plants.

**Methods:** We used analyses of diversity and endemism for 10% of the *campos rupestres flora*. We obtained distribution data from online databases, and phylogenetic hypotheses from the literature. With these datasets, we estimated alpha and beta metrics of taxonomic and phylogenetic diversity, and conducted categorical analyses of neo- and paleo-endemism.

**Results:** Phylogenetic overdispersion predominates in the *campos rupestres*. However, this general pattern is permeated by both lineage- and site-specific phylogenetic clustering, suggesting that recent diversification events depend on particular regional conditions and on the overall maintenance of old lineages. Although endemism patterns vary among different *campos rupestres* sites, paleo-endemism is widespread and particularly prominent where phylogenetic overdispersion is evident. Moreover, phylogenetic composition indicates variable past spatial connections across different sites, taxonomic composition is highly geographically structured and seems to be influenced by the vegetation surrounding the *campos rupestres* and/or by abiotic conditions.

**Main conclusions:** Our results reinforce the idiosyncratic nature of diversification patterns in ancient mountains and suggest that old, climatically buffered, infertile montane ecosystems not only include both relatively old and recent lineages, but that recent diversification is lineage and spatially dependent.

## 1 Introduction

Despite covering a strikingly small area of the globe, mountains house most of terrestrial biodiversity (Myers, Mittermeier, Mittermeier, Da Fonseca & Kent, 2000; Rafiqpoor, Kier, & Kreft, 2005; Kreft & Jetz, 2007; Rahbek et al., 2019). Their island-like nature and heterogeneous landscape promote diversification at multiple scales (Hughes & Eastwood, 2006; Drummond, Eastwood, Miotto & Hughes, 2012; Hughes & Atchison, 2015; Cortés, Garzón, Valencia & Madriñán, 2018; Nürk, Atchison, & Hughes, 2019; García-Rodríguez et al., 2021). However, the diversity of different mountains may have distinct origins. For example, new environments and geographic barriers that emerged during the Andes uplift led to fast and recent species diversification (Hughes & Eastwood, 2006; Särkinen, Pennington, Lavin, Simon & Hughes, 2012; Hughes, Pennington, & Antonelli, 2013). On the other hand, the long-term climatic and topographic stability of ancient mountains may have favoured the persistence of old lineages (Berry & Riina, 2005; Warren & Hawkins, 2006; Barbosa, Fernandes, & Sanchez-Azofeifa, 2015). Although these and other examples suggest that the evolution of montane diversity was intricate (e.g. Hughes & Atchison, 2015; Badgley et al., 2017; Antonelli et al., 2018), there is an ongoing debate over the main processes behind the origin of a hyper-diverse montane vegetation in Eastern South America (Silveira et al., 2016), the *campos rupestres*.

The *campos rupestres* are one of the richest and most endemic floras in the tropics (BFG, 2015). With more than 5000 plant species, it is similar to or almost twice as diverse as some biodiversity hotspots, such as Japan (5600 species), the Horn of Africa (5000 species), and New Zealand (2300 species) (Mittermeier, Turner, Larsen, Brooks & Gascon, 2011; Silveira et al., 2016). This floristic diversity is also spatially limited to an area smaller than Ireland or less than 1% of the Brazilian terrestrial territory (Silveira et al., 2016). *Campos rupestres* occur in old, climatically buffered, infertile landscapes (OCBIL; Silveira et al., 2016), over topographically heterogeneous and ancient quartzite, sandstone and ironstone formations ranging from 900 to over 2,000 m a.s.l. (Pedreira & De Waele, 2008; Fernandes, 2016). The vegetation mosaic of these areas, which include grasslands and rock-dwelling plant communities intermingled with gallery forest patches (for a detailed review, see Fernandes, 2016), occurs mostly in the Espinhaço Range, a mountain chain distributed along the Brazilian states of Minas Gerais and Bahia (Giulietti, Pirani, & Harley, 1997). Other disjunct sites, set apart by lowland vegetation are at the Brazilian Central Plateau (Goiás state and Distrito Federal) and at *Serra da Canastra* (western Minas Gerais state; Fig. 1; Silveira et al., 2016). The *campos rupestres* discontinuity is also reflected in species distribution, many of which are narrow and/or occur as small, disjunct populations (Giulietti et al., 1997; Echternacht, Trovó, & Takeo Sano, 2010; Echternacht, Trovó, Oliveira & Pirani, 2011; Lousada, Lovato, & Borba, 2013; Trovó, De Andrade, Sano, Ribeiro & Van den Berg, 2013).

**Figure. 1:**
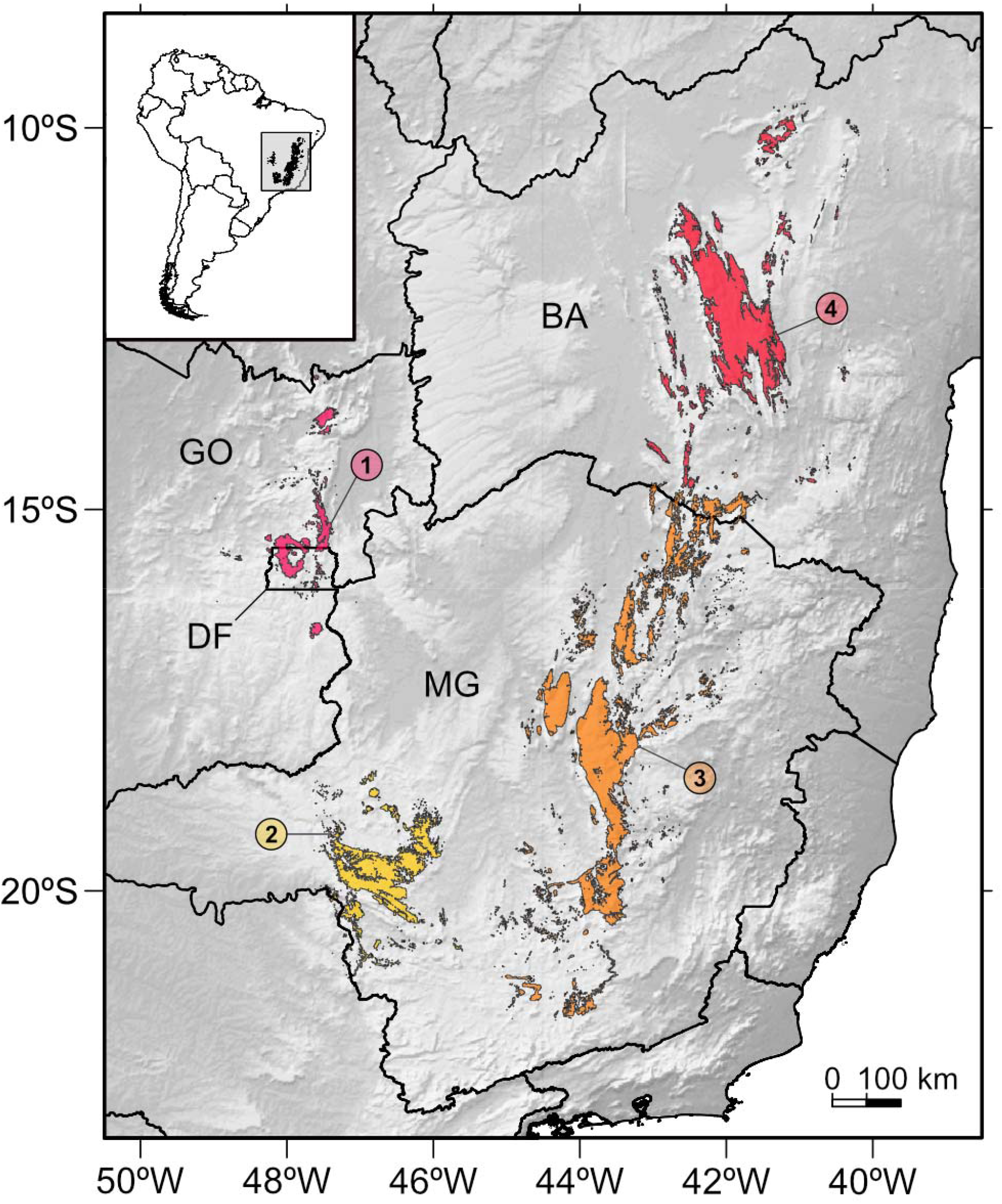
Main distribution of the *campos rupestres* within the Brazilian territory and across the Distrito Federal (DF) and the states of Bahia (BA), Minas Gerais (MG) and Goiás (GO). This vegetation complex is concentrated in the Espinhaço Range, but also occurs in the Brazilian Central Plateau and the *Serra da Canastra*. Here we considered the *campos rupestres* as divided into four different sections: the Southern and Northern Espinhaço, the Brazilian Central Plateau and the *Serra da Canastra*. Adapted from Silveira et al. (2016).

The distribution and diversity of *campos rupestres* plants have often been viewed as the result of a species-pump mechanism, which promotes recurrent bursts of diversification similar to the ones seen in the Andes (Harley, 1988; Giulietti et al., 1997; Flantua, O’dea, Onstein, Giraldo & Hooghiemstra, 2019). According to this hypothesis, recurrent climatic variations during the Tertiary and Quaternary (Adams, Maslin, & Thomas, 1999; Zachos, Dickens, & Zeebe, 2008) led to population expansion-contraction cycles and intermittent gene flow. In this context, isolation – sometimes together with hybridisation (e.g. Antonelli, Verola, Parisod & Gustafsson, 2010) – would promote local diversification (Giulietti et al., 1997; Vasconcelos et al., 2020). Indeed, shifts in the range of *campos rupestres* populations might have been common during the Pleistocene (Collevatti, Rabelo, & Vieira, 2009; Bonatelli et al., 2014; Barbosa, Fernandes & Sanchez-Azofeifa, 2015; Barres, Batalha-Filho, Schnadelbach & Roque, 2019), a period of high diversification (Souza et al., 2013; Trovó et al., 2013; Ribeiro, Rapini, Damascena & Van den Berg, 2014; Rando et al., 2016; Vasconcelos et al., 2020). Therefore, if these processes were as common as expected by the species-pump hypothesis, recently diverged lineages (i.e. neo-endemics) and signatures of recent diversification (e.g. phylogenetic clustering) should dominate *campos rupestres* communities.

However, similarly to other OCBILs, such as the Cape and southwestern Australia (Cowling et al., 2014), old communities should prevail in the *campos rupestres* due to the long-term climatic and topographical stability of these ancient mountains (Hopper, 2009). Although this prediction has been recently contested (Cortez et al., 2020), several evidences indicate that the *campos rupestres* may be one of the oldest modern day open vegetation in Tropical America, and include lineages diversified as early as the Oligocene/Miocene (Rapini, Berg, & Liede-Schumann, 2007; Antonelli et al., 2010; Gustafsson, Verola, & Antonelli, 2010; Hughes, Pennington & Antonelli, 2013; Zappi, Moro, Meagher & Nic Lughadha, 2017; Alcantara, Ree, & Mello-Silva, 2018). Thus, *campos rupestres* hyper-diversity would be, at least in part, due to the accumulation of old lineages.

Whichever processes were in action during formation of the flora of ancient mountains, their nature and/or intensity likely varied across space (Croizat, 1962; Nürk et al., 2020). However, little is known about how the *campos rupestres* high levels of topographical, climatic and edaphic heterogeneity (Giulietti et al., 1997; Schaefer, Cândido, Corrêa, Nunes & Arruda, 2016; Neves et al., 2018) shaped the distribution of old and/or recent lineages (but see Bitencourt & Rapini, 2013). Hence, to better understand the *campos rupestres*’ evolutionary history, it is necessary to evaluate how diversity is distributed across space.

Spatial analyses of phylogenetic diversity and endemism are reliable approaches to address this issue (e.g. Forest et al., 2007; Mishler et al., 2014; Heenan, Millar, Smissen, McGlone & Wilton, 2017; Scherson et al., 2017). Phylogenetic diversity (Faith, 1992) informs the proportion of recently and deeply diverged lineages in local communities (Webb, Ackerly, McPeek & Donoghue, 2002; Divya, Ramesh & Karanth, 2021), while the distinction between centres of neo- and paleo-endemism shows how these different types of lineages are distributed in space (Mishler et al., 2014). These patterns can be complemented by analyses of beta and phylobeta diversity (e.g. Jaccard, 1901; Lozupone & Knight, 2005), which point to past and present connections between areas, as well as to historical processes structuring communities (Graham & Fine, 2008).

Here we aimed to identify which evolutionary processes may have shaped the flora of an ancient Tropical mountain and how they are distributed across space. To do so, we assessed fine spatial patterns of phylogenetic diversity and endemism for over 10% of the *campos rupestres* flora. If predictions for the persistence of old lineages in ancient landscapes are met, patterns of phylogenetic overdispersion and paleo-endemism should prevail in the *campos rupestres*. On the other hand, widespread phylogenetic clustering and neo-endemism would indicate communities to be composed by recently diversified lineages, reinforcing the role of a species-pump mechanism in the diversification of altitudinal floras (e.g. Flantua et al., 2019; Vasconcelos et al., 2020). Moreover, a geographical organisation of the taxonomic and phylogenetic composition of the *campos rupestres* would indicate the biological connection of these mountains to be under topographical, climatic and edaphic constraints.

## 2 Methods

To answer our questions, we used 572 species from nine angiosperm groups for which well-sampled phylogenetic hypotheses are available: *Mimosa* L. (Leguminosae), *Diplusodon* Pohl (Lythraceae), *Comanthera* L.B. Sm. (Eriocaulaceae), Velloziaceae J. Agardh, Lychnophorinae Benth. (Asteraceae), *Paepalanthus* Mart. (Eriocaulaceae), *Calliandra* Benth. (Leguminosae), *Minaria* T.U.P. Konno (Apocynaceae) and Marcetieae M.J.R. Rocha, P.J.F. Guim. & Michelang. (Melastomataceae). Although this strategy could bias analyses towards the evolutionary history of sampled lineages, these groups are well-represented in the *campos rupestres*, their life forms range from herbs to trees and they comprise both monocots and eudicots. They are also mostly restricted to grasslands and rocky outcrops, commonly not occurring in the gallery forest patches dispersed across the *campos rupestres* area. Therefore, they are an adequate proxy for general *campos rupestres* diversity patterns.

Our analyses were divided into two parts: (1) data compilation and cleaning, and (2) spatial analyses. Below we present a general workflow for each step, all of which were conducted in the R statistical computing environment (R Core Team, 2013). For details about each analysis, see the Supporting Information (Appendix 1).

### 2.1 Data compilation and cleaning

We retrieved occurrence data for our model groups from two complementary databases:

GBIF (GBIF.org, 2020) and speciesLink (CRIA, 2020). Data extracted from these databases were concatenated into a single data set to facilitate posterior treatment. To reduce data dimensionality, we removed all variables but the ones relevant for data cleaning and spatial analyses.

To clean our data, we first pruned duplicated records and removed records without identification at species level, without names of determiners or without vouchers. Then we followed the methods proposed by Magdalena et al. (2018), which consist of: (1) defining a coordinate reference system; (2) standardising names of geographic units; (3) selecting and evaluating records with geographic coordinates; (4) identifying records without geographic coordinates; and (5) inferring geographic coordinates based on municipalities or field notes. To avoid bias due to species misidentification, we also updated the taxonomy, corrected typos in species names and limited the data set to records identified by taxonomic specialists (authors of monographs, floras or taxonomic revisions on the family of the focal group).

Finally, we removed all records whose coordinates fell outside the *campos rupestres*’ distribution (Silveira et al., 2016).

### 2.2 Spatial analyses

To understand the taxonomic and phylogenetic composition and structure of plant communities in the *campos rupestres*, we calculated the following metrics: species richness and phylogenetic diversity (PD; Faith, 1992), Jaccard distance (Jaccard, 1901) and UniFrac (Lozupone & Knight, 2005). We also conducted categorical analyses of neo- and paleo-endemism (CANAPE; Mishler et al., 2014) to assess the distribution of neo- and paleo-endemic lineages.

Since part of our analyses rely on phylogenetic trees, we obtained the most recent phylogenetic hypothesis for each group (Echternacht et al., 2014; Loeuille, Semir, Lohmann & Pirani, 2015; Alcantara et al., 2018; Inglis & Cavalcanti, 2018; Andrino et al., 2020; Vasconcelos et al., 2020), all of which include at least 50% of their diversity. Trees and distribution matrices were both pruned to include only species present in both. Also, we scaled every tree by total tree length, making it possible to compare the results for different groups and with future studies. All analyses were conducted at species level.

#### 2.2.1 Spatial grids

Although it is hard to define the best resolution of the spatial grids needed for diversity analyses (Blackburn & Gaston, 2002), their configuration affects the results of biogeographical studies (Willis & Whittaker, 2002; Rahbek, 2005). Thus, we based grid cells’ size on redundancy values, which take sampling effort in consideration (Scherson et al., 2017). To do so, we first removed grid cells that had only one sample per species. Then we calculated redundancy for all cells at different sizes and assessed the relationship between grid cells’ size and redundancy (i.e., the median of all redundancy values at a given cell’s size). To investigate if the diversity of each cell could be biased by unequal sampling effort, we estimated asymptotic species richness (Chao et al., 2014), a measure of “true” diversity, for each taxonomic group and for all of them combined. The high correlation between observed and estimated diversity (R2 ≥ 0.82 for two groups, R^2^ ≥ 0.93 for the other groups, R^2^ = 0.99 for all groups combined, and p-values < 0.05 for all groups individually and combined) indicates our dataset is evenly sampled and can be used in subsequent analyses without any corrections (e.g. interpolation or extrapolation (Chao et al., 2014). See the supporting information (Appendix 1) for details and results of these analyses.

#### 2.2.2 Alpha diversity

We calculated two metrics of alpha diversity: richness and phylogenetic diversity (PD). For PD, we used the ‘picante’ package (Kembel et al., 2010) to sum the branch lengths connecting a given set of species to the root of a phylogenetic tree (Faith, 1992). We also tested the correlation between richness and PD with Spearman’s correlation tests (Spearman, 1904).

Because PD is often correlated with richness (Faith, 1992; Polasky, Csuti, Vossler & Meyers, 2001; Rodrigues & Gaston, 2002; Tôrres & Diniz-Filho, 2004; Forest et al., 2007; Scherson et al., 2017), decoupling these two metrics provides a powerful tool to assess community phylogenetic structure and, thus, offers valuable insights into macroecological and biogeographical processes (Webb et al., 2002). We did this by using the residuals of linear regression models between phylogenetic diversity and richness. Residual values above or below zero indicate PD higher or lower than expected based on richness, respectively (Forest et al., 2007; Brown et al., 2020; Colville et al., 2020).

Finally, to gain a broader perspective on the spatial distribution of alpha diversity, we calculated total richness and total PD for all groups occurring in each grid cell. Then, we calculated the residuals of each metric for all groups combined (Brown et al., 2020).

#### 2.2.3 Beta diversity

To assess communities’ structure both in the present as in the past (Wiens & Donoghue, 2004; Emerson & Gillespie, 2008; Graham & Fine, 2008), we conducted UPGMA analyses for each group using dissimilarity matrices based on Jaccard distance (beta diversity; Jaccard, 1901) and UniFrac (phylobeta diversity; Lozupone & Knight, 2005). As with alpha diversity metrics, we also performed an UPGMA analysis based on Jaccard distance for all groups combined simply by merging their distribution matrices. This approach was not possible for UniFrac, as it relies on pairwise calculations based on unique phylogenetic trees.

Before generating the consensus dendrograms (1000 replicates; consensus criterion of 0.5) for each group, we removed singletons (species whose occurrence is restricted to a single cell) in order to avoid noise. Then, to better visualise spatial structure, we subdivided each dendrogram using fusion level values (Borcard, Gillet, & Legendre, 2018), while attempting to maintain a consistent number of clusters between all groups. All beta diversity analyses were conducted with

‘recluster’ (Dapporto et al., 2013).

#### 2.2.4 CANAPE

The categorical analyses of neo- and paleo-endemism were conducted following the methods developed by Mishler et al. (2014), which involve prior calculation of phylogenetic endemism (PE; Rosauer, Laffan, Crisp, Donnellan & Cook, 2009) and relative phylogenetic endemism (RPE; Mishler et al., 2014). After calculating PE and RPE with the ‘PDcalc’ package (D. Nipperess; under development), we performed all CANAPE’s statistical tests with a custom code. CANAPE includes four endemism categories: neo-endemism (rare, younger lineages), paleo-endemism (rare, older lineages), mixed endemism (a mix of neo- and paleo-endemics), and super-endemism (a strong signal of mixed endemism). Please bear in mind that the distinction between paleo- and neo-endemism is not absolute, but relative to the age of sampled lineages. In our analyses, we found some grid cells that could not be classified in any of CANAPE’s categories, as they do not meet the criteria for all its statistical tests. Thus, we defined their endemism as “uncertain”. See Supporting Information for details about this analysis (Appendix 1).

Because we did not include occurrence data from areas surrounding the *campos rupestres* adjacent areas, our endemism analyses could be biased. However, CANAPE relies on the range of phylogenetic branches, not that of terminals. Also, most *campos rupestres* species are restricted to small areas within it (e.g. Rapini, 2010; Echternacht et al., 2011; Bitencourt & Rapini, 2013; Trovó et al., 2013). Thus, we believe our analyses were robust enough to assess the distribution of different types of endemism within the *campos rupestres*’ range. Nevertheless, including the surrounding vegetation in future studies will certainly improve knowledge on these patterns.

## 3 Results

### 3.1 Redundancy

Redundancy values for all groups steadily increase together with grid cells’ size until reaching a plateau around 0.9 (Fig. S2.1). However, a value of 0.9 for redundancy defines cells’ sizes too large to recover variations in spatial patterns. Hence, aiming for a balance between resolution and redundancy, we used cells of 0.6 per 0.6 decimal degrees (roughly 66 km per 66 km), whose redundancy varies between 0.6 and 0.8.

### 3.2 Richness and phylogenetic diversity

Richness in the *campos rupestres* is concentrated in the Southern Espinhaço, where some cells include more than 200 species, and in the Brazilian Central Plateau (Figs. 2a and S2.2). The richness peak in the national capital could be a bias due to automatic georreferencing using countries’ capitals (Zizka et al., 2019) or higher collecting effort in this area, which harbours several herbaria and research institutions. However, as some of our model groups are very diverse in this area (*Mimosa* and *Diplusodon*; Barneby, 1991; Cavalcanti, 1995; Cavalcanti, 2007) and our dataset is relatively evenly sampled, we decided to keep records for this particular cell to avoid underestimation of *campos rupestres* diversity in the Brazilian Central Plateau.

**Figure 2:**
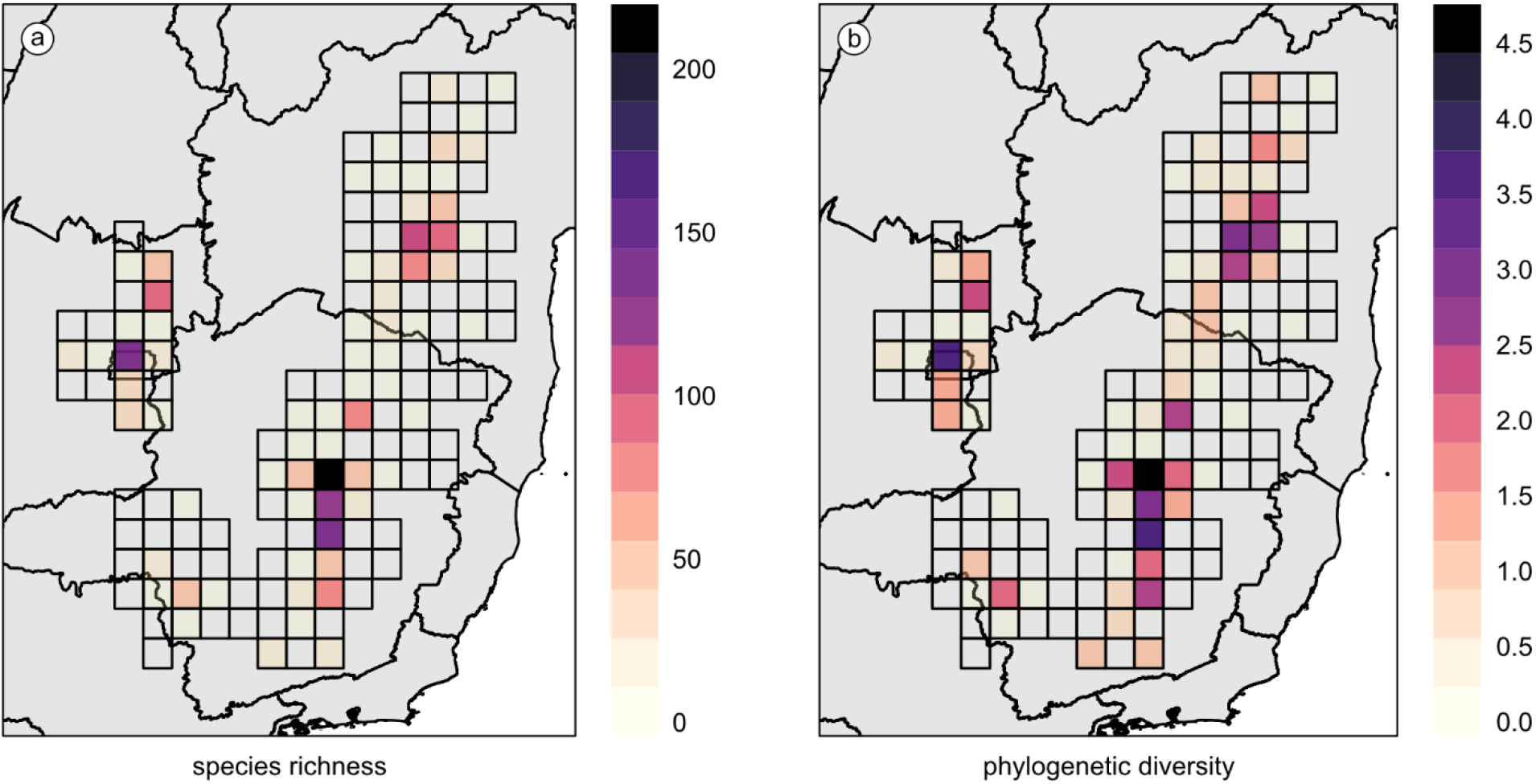
Combined results of (a) species richness and (b) phylogenetic diversity for all groups. Phylogenetic trees were scaled by total tree length so phylogenetic diversity results could be combined. Transparent grid cells indicate absent data

As expected (Faith, 1992; Polasky et al., 2001; Rodrigues & Gaston, 2002; Tôrres & Diniz-Filho, 2004; Forest et al., 2007; Scherson et al., 2017), PD follows richness distribution (Fig. S2.3), and highest PD values occur in the Southern Espinhaço, followed by the Brazilian Central Plateau and the Northern Espinhaço (Figs. 2b and S2.4). Although we noticed some individual groups may dominate and skew the residuals of the regression between total richness and total PD (Fig. S2.5), the combined residuals of all groups present a more accurate general view of the phylogenetic structure of the *campos rupestres* communities.

The PD of a large proportion of *campos rupestres* communities follow what is expected by richness (50% of residual values range from -0.2 to 0.2), particularly in the Brazilian Central Plateau. Nonetheless, the residual values show that communities in which PD is higher than expected by richness (phylogenetic overdispersion; positive residuals) are more common than communities in which PD is lower than the expected (phylogenetic clustering; negative residuals), more so in the Espinhaço Range and *Serra da Canastra* (Fig. 3). Although a single Southern Espinhaço site has an extremely low residual value, phylogenetic overdispersion is usually stronger than phylogenetic clustering (Fig. 3). The highest levels of phylogenetic overdispersion occur in the Northern Espinhaço, particularly when individual groups are considered.

**Figure 3:**
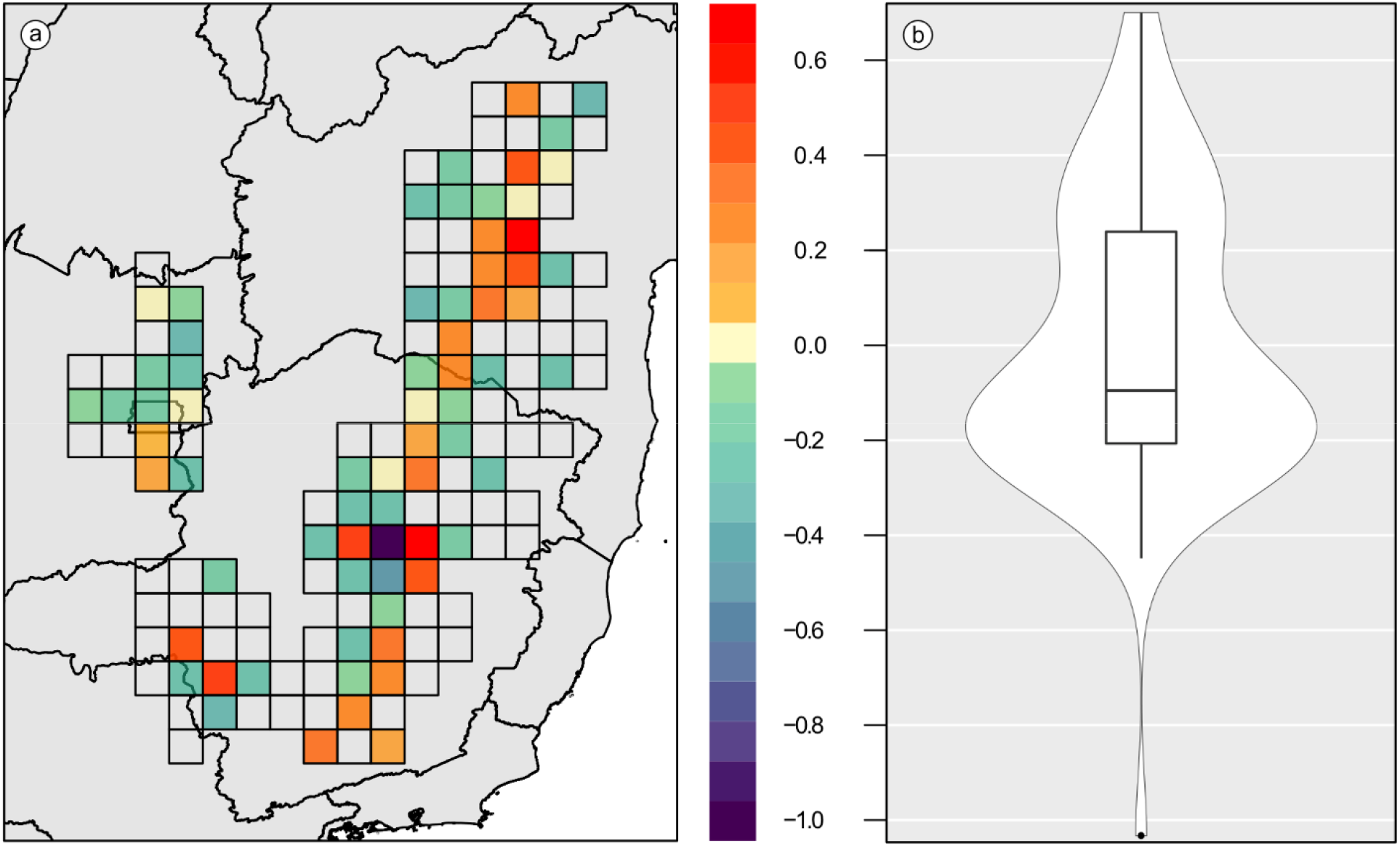
Spatial distribution of residuals from linear regressions between species richness and phylogenetic diversity across the *campos rupestres* for all groups combined (a). Positive and negative residuals respectively indicate phylogenetic overdispersion and clustering (i.e. communities in which phylogenetic diversity is higher or lower than expected based on richness). Transparent grid cells indicate absent data). The violin boxplot (b) depict the distribution of residuals along the total range.

Residual values for each focal group (Fig. S2.6) agree with a general pattern of phylogenetic overdispersion, but also show variations in the occurrence of phylogenetic clustering. For example, phylogenetic clustered communities not clearly seen in the joint analysis occur in the Brazilian Central Plateau (*Mimosa* and *Diplusodon*), in the Southern Espinhaço (*Comanthera* and *Minaria*), in the Northern Espinhaço (Marcetieae and *Calliandra*), and across the whole Espinhaço Range (*Paepalanthus*).

### 3.3 CANAPE

Although the distribution of endemism categories varies among individual groups, CANAPE results indicate that paleo-endemic lineages predominate in the Northern Espinhaço, while other *campos rupestres* areas include a mix of paleo- and neo-endemism (Fig. 4; see also Fig. S2.7 for detailed information on each group). *Mimosa*, Velloziaceae and Lychnophorinae are paleoendemic to the Northern Espinhaço, which also includes a case of mixed endemism for *Comanthera*. All endemism categories occur in the Southern Espinhaço: on its northern part, paleo-, super and mixed endemism are seen for *Paepalanthus, Comanthera* and Velloziaceae, whereas super, mixed, paleo- and neo-endemism of *Paepalanthus, Comanthera, Calliandra*, and *Minaria* all occur on its northern part. Finally, mixed endemism for *Diplusodon* and Velloziacae defines the *Serra da Canastra*, while neo-, paleo- and super endemism characterise the Brazilian Central Plateau (*Mimosa, Diplusodon, Comanthera* and Marcetieae).

**Figure 4:**
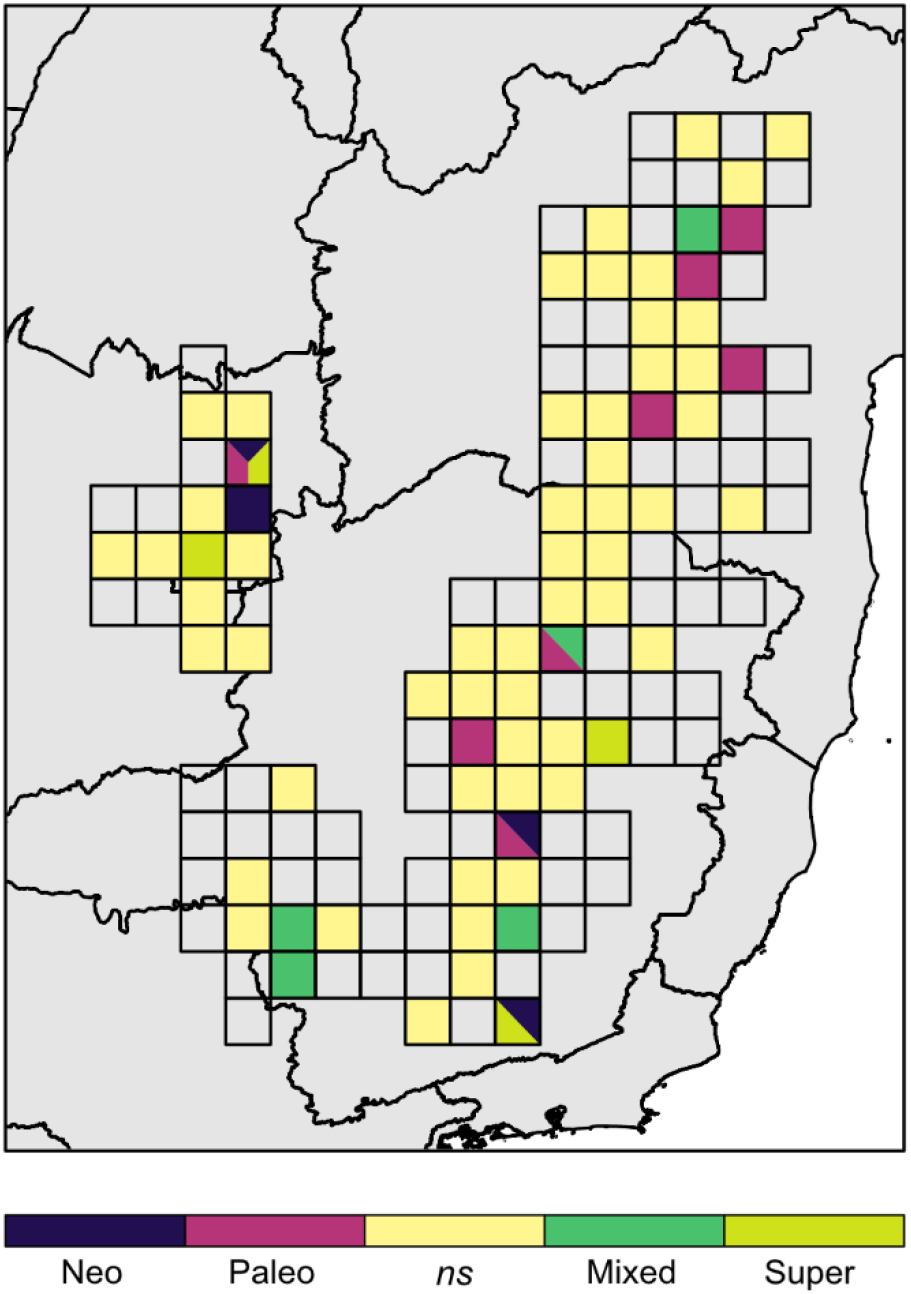
CANAPE results of all groups combined. Neo-endemic cells contain a predominance of short, rare phylogenetic branches; paleo-endemic cells contain a predominance of long, rare branches; and cells with mixed endemism contain a mixture of both. Mixed endemism can be further categorised as super endemism in a cell in which it is statistically significant at a lower confidence interval (*α* ≤ 0.001). Combining the results for all groups results in some cells harbouring multiple endemism types. Transparent grid cells indicate absent data.

It is important to note that we were not able to categorise a few grid cells for *Comanthera*. This is likely due to low statistical power: *Comanthera* is a small genus (Echternacht et al., 2014), and the number of species assessed in this work was further reduced so that the phylogenetic tree could match the distribution matrix (see Methods). Thus, low sampling may have negatively affected the permutations performed during CANAPE analyses.

### 3.4 Jaccard distance and UniFrac

UPGMA analyses based on Jaccard distance (Fig. 5) showed that plant communities are geographically structured. Although the limits of these geographical patterns might have changed over time, the Northern Espinhaço flora stands in contrast with the area encompassing the Brazilian Central Plateau, the *Serra da Canastra* and the Southern Espinhaço Range (hereafter referred to as the Central Plateau-Southern Espinhaço arch).

**Figure 5:**
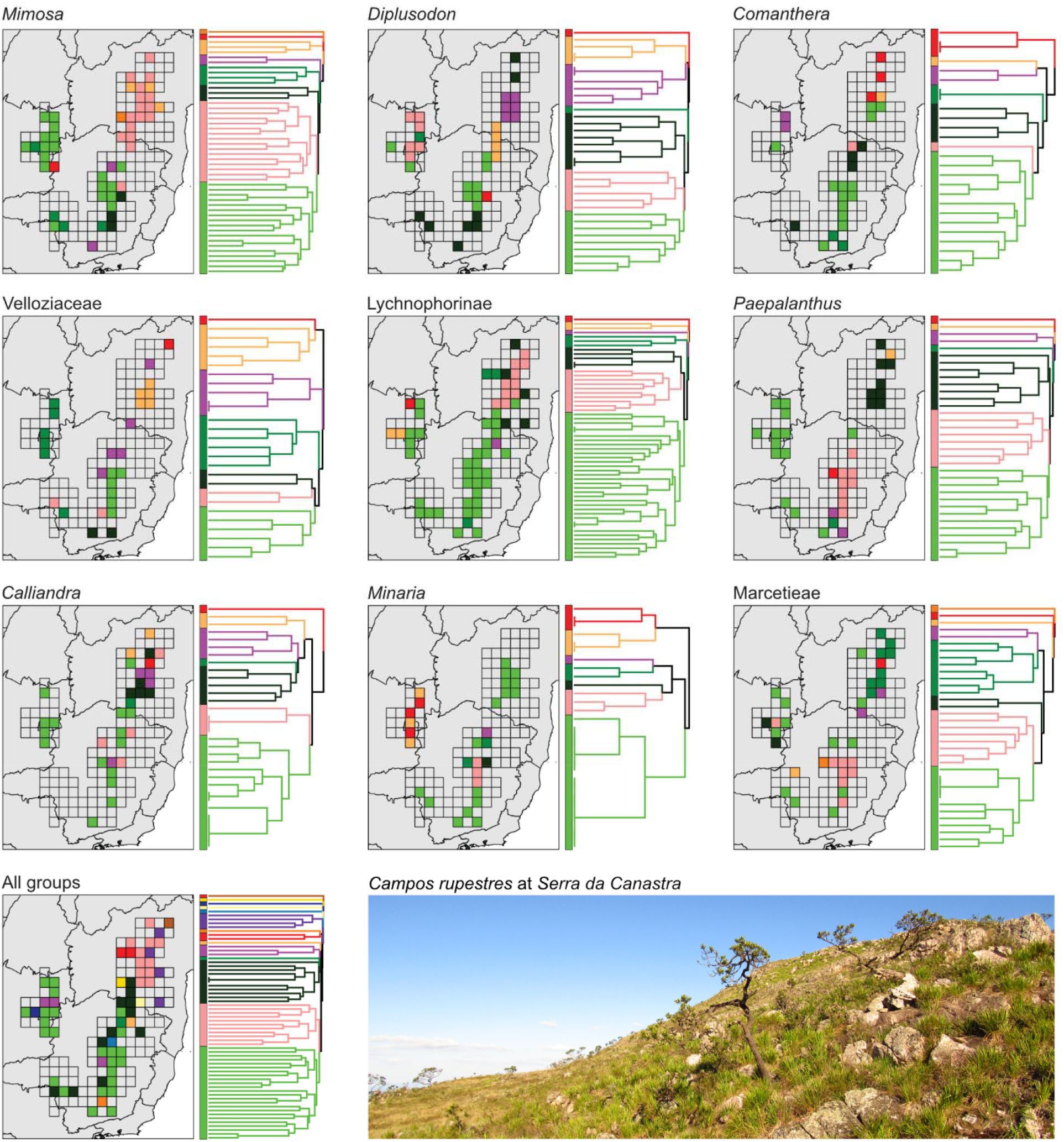
Hierarchical clustering of *campos rupestres* communities based on their taxonomic composition. Colours and cells in the map correspond to colours and terminals in the dendrogram, respectively. The closer the terminals are, more taxonomic identity the corresponding cells share. Transparent grid cells indicate absent data. See supporting information for bootstrap values.

The remarkable contrast between the taxonomic composition of the Northern Espinhaço and that of other *campos rupestres* is evident for all groups combined and also for each taxon individually, attesting to the consistency of this pattern (Figure 5). The floristic cohesiveness between the Brazilian Central Plateau, the *Serra da Canastra* and the Southern Espinhaço is also very strong. Even in the few cases in which the Brazilian Central Plateau and/or the Southern Espinhaço appear as taxonomically unique (e.g. *Diplusodon* and *Paepalanthus*), these regions are nested within a larger Central Plateau-Southern Espinhaço group.

A divide between the Central Plateau-Southern Espinhaço and the Northern Espinhaço is also seen in the phylogenetic composition of *campos rupestres* communities, particularly in *Mimosa, Diplusodon*, Velloziaceae, *Paepalanthus* and Marcetieae, but with fuzzier geographic limits (Fig. 6). For example, the results for *Mimosa* extend the limits of a Central Plateau-Southern Espinhaço’s cluster further north, while a Northern Espinhaço’s cluster includes close areas in the Southern Espinhaço.

**Figure 6:**
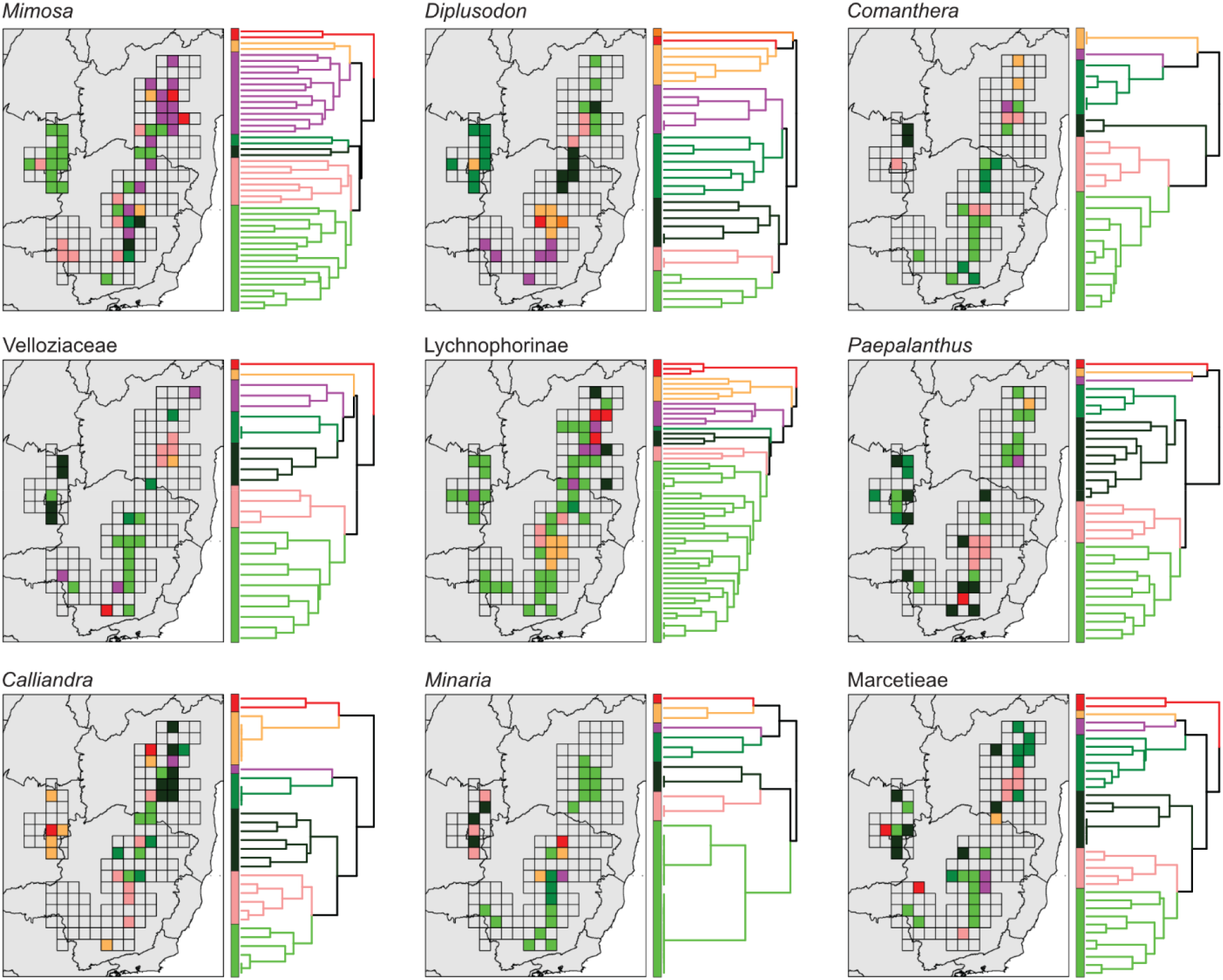
Hierarchical clustering of *campos rupestres* communities based on their phylogenetic composition. Colours and cells in the map correspond to colours and terminals in the dendrogram, respectively. The closer the terminals are, more phylogenetic identity the corresponding cells share. Transparent grid cells indicate absent data. See supporting information for bootstrap values.

There are exceptions to this general geographic structure. For example, in *Calliandra* the Brazilian Central Plateau appears as the most evolutionary contrasting region, while the Southern and Northern Espinhaço form a single, cohesive region. For *Minaria*, the Espinhaço Range and the *Serra da Canastra* form a cluster that contrasts to the Brazilian Central Plateau, even though both regions are nested into a widespread cluster. Moreover, the central Southern Espinhaço frequently appears as an evolutionary unique area, as seen for *Mimosa, Diplusodon*, Lychnophorinae, and *Minaria*.

## 4 Discussion

Aiming to understand the evolutionary history of ancient tropical mountains, here we investigated spatial patterns of richness, phylogenetic diversity and endemism for the *campos rupestres* mega-diverse flora. We also assessed phylogenetic and taxonomic differences between communities using beta and phylobeta diversity metrics. Although particular results reinforce that the *campos rupestres* are a complex assemblage of different evolutionary histories, these are not evenly distributed across space. Below we discuss the relationship between these spatial patterns and what processes might have shaped them.

### 4.1 Spatial distribution of recent and old lineages

Richness and phylogenetic diversity are not evenly distributed in the *campos rupestres*, but concentrated in the Brazilian Central Plateau and the Southern Espinhaço. Decoupling these two metrics from each other reveals different aspects of community composition. The combined residuals of all groups indicate that phylogenetic overdispersion is more common than phylogenetic clustering, even though PD follows what is expected by richness in a large number of sites (ca. 50% of grid cells). The particular predominance of expected PD values in the Brazilian Central Plateau, however, must be seen with caution, as there may be a bias against groups whose communities are phylogenetically clustered in this region (e.g. *Mimosa* and *Diplusodon*; Fig. S2.5). Regarding endemism patterns, paleo-endemism predominates in the Northern Espinhaço, while the Southern Espinhaço, *Serra da Canastra* and Brazilian Central Plateau floras (Central Plateau-Southern Espinhaço arch) include a mixture of paleo- and neo-endemic lineages.

Widespread phylogenetic overdispersion indicates the predominance of relatively older (deeply diverged) lineages (Lososová et al., 2015; Colville et al., 2020; Divya et al., 2021). CANAPE reinforces this by showing that endemics commonly belong to old lineages (paleo-endemism). This pattern is especially prominent in the Northern Espinhaço, where both paleo-endemism and phylogenetic overdispersion predominate. Contrasting with the Northern Espinhaço, the Central Plateau-Southern Espinhaço arch includes more areas of phylogenetic clustering, with a single Southern Espinhaço site showing the lowest residual value (close to -1), and a mix of paleo- and neo-endemism or even well-marked neo-endemism. This pattern is clearer in analyses of individual taxa, especially in the Brazilian Central Plateau, where neo-endemism for *Mimosa* and *Diplusodon* overlaps with phylogenetically clustered communities. These results show that the Central Plateau-Southern Espinhaço arch flora includes patches of relatively young (short branches) and closely related lineages (Webb et al., 2002; Divya et al., 2021), as previously indicated by phylogenetic and diversification analyses (Trovó et al., 2013; Bonatelli et al., 2014; Rando et al., 2016; Vasconcelos et al., 2020).

Taken together, these patterns indicate that recent diversification is spatially restricted, lineage-dependent and embedded in a widespread and relatively older flora (Figures 3 and 4). As expected (Hopper, Silveira, & Fiedler, 2016; Silveira et al., 2016; Hopper, Lambers, Silveira & Fiedler, 2021), long-term climatic and topographical stability (Pedreira & De Waele, 2008; Silveira et al., 2016) may have favoured the persistence of old lineages, promoting an accumulation of species over time similar to that seen in the Cape and southwestern Australia (Cowling et al., 2014). Such a reservoir of older lineages was likely the raw material for bursts of diversification, which could have been driven by the reassembly of ancient alleles (Richards et al., 2021), for example. Moreover, high heterogeneous topography, specific edaphic conditions, and particular ecophysiological requirements (Giulietti et al., 1997; Fernandes, 2016; Oliveira et al., 2016, Schaefer et al., 2016) likely constrained plant diversification across the *campos rupestres*. Thus, although probably present (e.g. (Antonelli et al., 2010; Bonatelli et al., 2014; Barbosa et al., 2015; Barres et al., 2019), climatic driven species-pumps apparently were not the main process shaping floristic diversity in the *campos rupestres* and other ancient mountains (e.g. Onstein & Linder, 2016).

A matrix of phylogenetic overdispersed communities could have been assembled by the free flow of multiple unrelated lineages across the vegetation surrounding the *campos rupestres*, as predicted by a model invoking changes in niche breadth as the precursors of lineage range expansion and diversification (Rapini, Bitencourt, Luebert & Cardoso, 2020). However, processes other than the balance between adaptive radiations and extinctions after drift invoked by this model could have shaped *campos rupestres* diversity. Hybridisation, for example, although relatively rare in OCBILs (Hopper, 2018), is widespread in angiosperms (Rieseberg & Willis, 2007), not uncommon in the *campos rupestres* (e.g., Borba & Semir, 1998; Ribeiro et al., 2008; Yamagishi-Costa & Forni-Martins, 2009), and often linked to macroevolutionary diversification events (Rieseberg & Willis, 2007). Also, species already well-established in the *campos rupestres* were likely less prone to extinction than species of similar ecology generated during bursts of recent diversification (McPeek, 2007). The role of such ecologically similar and transient species in the *campos rupestres* flora is reinforced by the occurrence of phylogenetic clustering in species-rich sites (McPeek, 2007), shown here for individual taxa, as well as by the overall high proportion of short branches seen for the Central Plateau–Southern Espinhaço arch (Cortez et al., 2020). Repetition of these localised birth and death events over time could produce the spatter of recent diversification within a matrix of relatively older lineages we observed here. Why local diversification is more common in the Central Plateau–Southern Espinhaço arch than in the Northern Espinhaço can be better discussed in the context of beta and phylobeta diversity analyses.

### 4.2 Present and past geographical structure of plant communities

As expected for island-like systems (e.g. Garcia-Verdugo, Forrest, Balaguer, Fay & Vargas, 2010; Kubota, Hirao, Fujii & Murakami, 2011; Carstensen et al., 2012), beta diversity analyses show a geographic structuring of the *campos rupestres* flora. Although the Southern Espinhaço and the Brazilian Central Plateau differ in species composition, together with the *Serra da Canastra* they form a cohesive group (the Central Plateau-Southern Espinhaço arch), which stands in taxonomic and phylogenetic contrast to the Northern Espinhaço. The limits between these two major areas, however, apparently varied across time.

A floristic dissimilarity between the Northern Espinhaço and the Southern Espinhaço has been previously stressed (Harley, 1988; Rapini, 2010; Bitencourt & Rapini, 2013; Trovó et al., 2013; Neves et al., 2018) and attributed to the role of the lowland between the two portions of the Espinhaço Range as a barrier to migration (Harley, 1988; Bitencourt & Rapini, 2013). However, even though *campos rupestres* species are known for their limited dispersal potential (Giulietti et al., 1997; Conceição et al., 2016; Arruda et al., 2020), the connections between distant areas belonging to the Central Plateau-Southern Espinhaço arch questions the role, or at least the strength, of such geographical barriers. Indeed, changes in community composition across the *campos rupestres* may be better predicted by environmental conditions (Neves et al., 2018): while the Northern Espinhaço is relatively less fragmented and located in the driest extreme of the Brazilian precipitation gradient, the Central Plateau-Southern Espinhaço arch includes highly fragmented sites and lies in a relatively moister environment (Bitencourt & Rapini, 2013; Neves et al., 2018). At the same time, the propensity of closer sites to share lineages indicates that geographic proximity may have also influenced the assembly of *campos rupestres* communities. Although the role of distance has been dismissed for the *campos rupestres* woody flora (Neves et al., 2018), our analyses including both woody and herbaceous taxa indicate it should be further tested.

Differences in the type of vegetation surrounding the *campos rupestres*, which have a remarkable influence on its woody flora (Neves et al., 2018; Assunção-Silva & Assis, 2021), probably also shaped the distribution of species composition. While the Central Plateau-Southern Espinhaço arch is chiefly embedded in a savanna-like vegetation (the Brazilian *cerrado*), the Northern Espinhaço is predominantly surrounded by a massive seasonally dry tropical forest, the *caatinga* (Conceição et al., 2016; Neves et al., 2018). One example of the role of the surrounding vegetation comes from *Calliandra*, in which a lineage from the *caatinga* radiated into a clade mostly endemic to the Northern Espinhaço (Souza et al., 2013). Similarly, radiations in the Cape usually occurred after transitions from forest to fynbos (Onstein, Carter, Xing & Linder, 2014), reinforcing the idea that adjacent environments might have played a major role in the assembly of mega-diverse flora in ancient mountains. Different processes could explain how lineages move from one vegetation to the another, including exaptations (*sensu* Gould & Vrba, 1982), the previously mentioned changes in niche breadth (Rapini et al., 2020), or sharing of similar environmental filters. Although these processes are not mutually exclusive, if the *campos rupestres* distribution area was not affected by Pleistocene climatic fluctuations (Rapini et al., 2020), lineage migration between disjunct sites must have occurred through the vegetation surrounding the *campos rupestres*. Thus, a scenario in which the *cerrado* chiefly exchanged lineages with the Central Plateau-Southern Espinhaço arch, while the Northern Espinhaço interacted mainly with the *caatinga* flora helps to explain the division of the *campos rupestres* into these two distinct floristic groups.

The putative influence of the surrounding vegetation may also explain phylogenetic diversity and endemism patterns, particularly the predominance of neo-endemism and phylogenetic clustering in the Central Plateau-Southern Espinhaço Arch. Because the *caatinga* is also an ancient and climatically stable habitat (Werneck, Costa, Colli, Prado & Sites Jr, 2011; Queiroz, Cardoso, Fernandes & Moro, 2017), a higher proportion of paleo-endemic and deeply diverged lineages are expected to colonise the Northern Espinhaço. On the other hand, the effects of Pleistocene climatic fluctuations over the more recently assembled *cerrado* (Behling, 2002; Safford, 2007; Simon et al., 2009; Werneck, Nogueira, Colli, Sites Jr & Costa, 2012) could have promoted recent lineage- and site-specific diversification in the Central Plateau-Southern Espinhaço arch, especially if morphological innovations matched climatic shifts followed by the emergence of ecological opportunities (Onstein et al., 2014). Nonetheless, it is important to note that the Northern Espinhaço and the Central Plateau-Southern Espinhaço arch are not completely evolutionary independent, as shown by the variations in their limits seen for different lineages and across relative time (Fig. 6). This indicates that, even though some lineages were unable to disperse (possibly due to intrinsic limitations, as suggested by a few evolutionary unique areas; Fig. 6), disjunct sites were probably connected multiple times and by different lineages in the past. Whether expansion of the *campos rupestres* area (e.g. Antonelli et al., 2010), niche shifts to the surrounding vegetation (Rapini et al., 2020), or other processes, such as phenotypic plasticity, promoted lineage dispersion is still debatable. Regardless of these uncertainties, it is clear that studies incorporating multiple aspects, such as abiotic, biotic, and clade-specific features of the *campos rupestres* and their surrounding floras may help to elucidate how historical connections between disjunct ancient mountains were possible.

## 5 Conclusion

Competing evolutionary processes have been evoked to explain the origins and diversity of both young and ancient mountains across the world (Hopper, 2009; Onstein et al., 2014; Flantua et al., 2019; Rapini et al., 2020; Vasconcelos et al., 2020), including long-term lineage persistence (Hopper, 2009), changes in niche breadth followed by range expansion (Rapini et al., 2020), and local diversification driven by climatic shifts (Vasconcelos et al., 2020) among others. However, even though evolution is strongly dependent on space (Croizat, 1962; Nürk et al., 2020), species and lineage distribution are often neglected when assessing the role of these processes in the assembly of mega-diverse floras, such as those of Eastern South American mountains, the *campos rupestres*. Here we aimed to assess the distribution of multiple evolutionary processes across the landscape of one of the richest floras in the world by combining spatial and phylogenetic data.

We show that the *campos rupestres* are split into two large areas of cohesive floristic diversity: the Central Plateau-Southern Espinhaço arch and the Northern Espinhaço. Although likely variable in the past, the contrast between these regions is reinforced by patterns of phylogenetic diversity and endemism. While phylogenetic overdispersion and paleo-endemism are dominant in the Northern Espinhaço, phylogenetic clustering and neo-endemism are mostly restricted to the Central Plateau-Southern Espinhaço arch, with very few exceptions. This is not only in agreement with patterns of floristic composition, but also with the distribution of environmental conditions and surrounding vegetation (Neves et al., 2018), which might have played an important role in the assembly of old, climatically buffered, infertile vegetations (Hopper, 2009; Onstein et al., 2014; Hopper et al., 2016).

Overall, this species-rich, highly endemic vegetation is mainly composed by relatively older lineages, spattered with patches of recently diversified lineages. Therefore, our spatial approach reconciles the apparent conflicting evidences for recent and fast diversification and the persistence of old lineages in ancient, climatic buffered, infertile landscapes, such as the Greater Cape Floristic Region (Linder, 2008; Cowling et al., 2014) and the *campos rupestres* (Simon et al., 2009; Bitencourt & Rapini, 2013; Hughes et al., 2013; Souza et al., 2013; Trovó et al., 2013; Bonatelli et al., 2014; Silveira et al., 2016). Nonetheless, our work shows that recent diversification events in long-term stable mountains can be both lineage- and site-dependent and may rely on the overall maintenance of relatively older lineages. These findings emphasise the idiosyncratic nature of diversification events and highlight the importance of considering both the spatial component and intrinsic lineage aspects when investigating diversity-related questions, as space clearly affected the balance between different evolutionary processes behind the astonishing diversity of ancient mountains.

## Acknowledgments

We thank Fernando Silveira, Rosa Scherson, Suzette Flantua, Rafael Izbicki, Renske Onstein, Kaori Nagata, two anonymous reviewers and the editors for their criticisms and suggestions. YB was supported by the Coordenação de Aperfeiçoamento de Pessoal de Nível Superior - Brasil (CAPES) - Finance Code 001. No permits were needed to carry this work.

## Data availability statement

The data that support the findings of this study are openly available at https://zenodo.org/record/7226249 (DOI: 10.5281/zenodo.7226249).

## Biosketch

**Yago Barros-Souza** is interested in the biogeography of neotropical vegetations and the role of morphological traits in plant evolution. This work was carried out as his master’s thesis at the Universidade de São Paulo, Brazil. **Leonardo M. Borges** studies the taxonomy and evolution of *Mimosa* and related genera in the legume family. He is also interested in the theory and practice of ontology, homology, and morphological disparity of plants. Except for data collection and analyses, led by YB, both authors contributed equally to this work.

Editor: Carina Hoorn

