## Supporting Information for "Spatial and lineage dependent processes underpin floristic assembly in the megadiverse Eastern South American mountains"

### Appendix S1: Methods

#### S1.1 Data compilation and cleaning

We retrieved occurrence data in the whole Brazilian territory for our model groups from two databases, specifically GBIF (GBIF.org, 2020) and speciesLink (CRIA, 2020). Occurrence data from these databases were then concatenated into a single data set for each group in order to facilitate posterior treatment. To reduce data dimensionality, we removed all variables but 15, which included information relevant for data cleaning and spatial analyses. These were: institution code, collection code, collection catalogue number, generic epithet, specific epithet, infraspecific epithet, original database name, names of determiners, record number, basis of record, collectors' names, federative unity of occurrence, municipality of occurrence, field notes, latitude and longitude. We also assigned each entry with an unique identifier number.

To clean our data, we followed the methods proposed by Magdalena et al. (2018), with some changes included. The cleaning procedure was divided into five steps: (1) defining a coordinate reference system; (2) standardising names of determiners, filtering by taxonomic specialists and checking for typos and taxonomic synonyms; (3) selecting and evaluating records with geographic coordinates; (4) identifying records without geographic coordinates; and (5) standardising names of geographic units and inferring geographic coordinates based on municipalities or field notes. These steps were preceded by a pre-refinement routine.

#### S1.2 Pre-refinement

Although speciesLink and GBIF have independent data, they also share identical records that would be duplicated in the merged data set. In order to remove duplicated information, we kept only one copy of records whose values in the following attributes were identical: institution code, collection code and collection catalogue number. We did so because redundancy analysis (see below) relies on the ratio between richness and sampling. Also, a reduced number of records facilitate cleaning procedures.

Next, we removed records without identification at species level, without names of determiners or without vouchers (field observations). We also standardised federative unities names and kept only observations from federative unities covering the *campos rupestres* (i.e. Bahia, Minas Gerais, Goiás, Federal District, Mato Grosso, Paraíba and Pernambuco; Silveira et al. 2016). We chose to maintain records with no information on federative unity of occurrence in order to avoid losing data with information on geographic coordinates, municipality of occurrence or field notes. These records would be cleaned in following steps.

#### S1.3 Coordinate reference system

We established the World Geodetic System (WGS84) as our coordinate reference system. Considering that WGS84 is the reference frame used by the Global Positioning System (GPS), we found it appropriate when dealing with coordinates generated by a GPS receiver device, which is the case for occurrence data retrieved from GBIF and speciesLink.

#### S1.4 Filtering by taxonomic specialists and checking for taxonomic synonyms

Taxonomically unreliable data is a source of bias in spatial diversity analyses (Goodwin et al., 2015). To avoid this issue, we filtered the data set by determiners' names, keeping only observations identified by taxonomic specialists. We defined a taxonomic specialist as an author of monographs, floras or taxonomic reviews on the family of the focal group. This step was preceded by a semi-automatic routine for standardisation of determiners' names based on Jaro-Winkler distances (Jaro, 1989). We manually checked determiners' names with a similarity of 70% or more with any of the taxonomic specialists names.

Also, to further improve the quality of our data, we checked for taxonomic synonyms and typos in species' names. We did this by generating a list with all species' names and manually evaluating them with the aid of Tropicos (Tropicos.org, 2020), The Plant List (The Plant List, 2020) and Reflora (Reflora, 2020). Synonyms were replaced by accepted names and typos were corrected. We removed records of species whose names were not found in any of the aforementioned platforms.

#### S1.5 Evaluation and inference of geographic information

After removing taxonomic dubious data, we focused on cleaning geographic information. This procedure was divided into two steps. First, we evaluated coordinate quality, and then we inferred georeference data for records lacking or with invalid coordinates.

Authenticity of geographic information was assessed with the 'CoordinateCleaner' package (Zizka et al., 2019). Records fitting one or more of the following criteria were considered invalid: (1) identical or plain zero values for latitude and longitude; (2) points falling in oceans; (3) geographic isolation (not applied to species with less than seven records); and (4) proximity to biodiversity institutions. 'CoordinateCleaner' also flags records near the centroid of a country's capital by default, but we did not consider them invalid because the Brazilian capital is an area of interest.

In order to infer coordinates for records without or with invalid georeferences, we used data on municipality of occurrence or field notes. To do so, we first standardised these two attributes by removing accents and special characters and capitalising major words. We then extracted a list of municipalities in which the *campos rupestres* occur by overlapping the Brazilian municipalities (IBGE, 2019) with the distribution of the *campos rupestres* according to Silveira et al. (2016). Next, we standardised this list following the same routine as for municipalities and field notes, filtered our data set and inferred coordinates for each record based on the centroid of its municipality. For records without this information, we used field notes, as these often mention the municipalities in which the records occur. Finally, we removed all records, including those evaluated in previous steps, whose coordinates fell outside the *campos rupestres*' distribution, making it ready for the spatial analyses and ensuring a data set comprised only by species occurring within the area of interest.

#### S1.6 Spatial grids and sampling evenness

The analyses of spatial patterns of diversity primarily depend on the establishment of a spatial grid cells' size. Although there is no way to objectively define the best resolution (Blackburn & Gaston, 2002), this step is fundamentally important, since grid resolution could affect the results of biogeographical studies (Rahbek, 2005; Willis & Whittaker, 2002). Thus, in order to select adequate grid cells' size, we used redundancy values, which assesses sampling effort by considering both richness and number of samples (1-[richness/(number of samples)]). A redundancy value close to one indicates good sampling; zero represents only one sample per taxa (Scherson et al., 2017).

Once we generated grid cells based on the distribution area of the *campos rupestres*, with sizes ranging from 0.1 per 0.1 DD to 5.0 per 5.0 DD, we calculated redundancy for all cells at different sizes and assessed the relationship between grid cells' size and redundancy (calculated as the median of all redundancy values at a given cell's size). Redundancy, although somehow arbitrary, aided us to find a good trade-off between grid cell' size and sampling. After choosing the best grid resolution, we removed grid cells that had only one sample per species, thus avoiding unreliable data. This eliminated cells located in federative unities in which the *campos rupestres* are less represented, such as Mato Grosso, Paraíba and Pernambuco.

Because sampling effort if often unevenly distributed (Boakes et al., 2010), results that rely on geographical occurrence may be biased towards areas where sampling effort is higher (e.g. areas that harbour research institutions and herbaria). Thus, to identify potential biases in sampling effort in our dataset, we estimated asymptotic species richness (estimated true diversity) using a protocol based on Hill numbers (Chao et al., 2014; Hsieh et al., 2016). Then, we ran linear models to understand if there was a mismatch between observed and estimated species richness. Observed and estimated species richness are highly correlated for each group individually (R2 not lower than 0.82 for two groups and higher than 0.93 for the others; p-value lower than 0.05) and when their diversity is combined (R2 = 0.99; p-value lower than 0.05). These results indicate sufficient sampling in the campos rupestres for the model groups used in our analyses. In this context, corrections based on interpolation/extrapolation are not required (see Figure S2.9).

#### S1.7 Alpha diversity

We calculated two metrics of alpha diversity: species richness and PD (Faith, 1992). We also tested the commonly found correlation between them (Faith, 1992; Forest et al., 2007; Polasky et al., 2001; Rodrigues & Gaston, 2002; Scherson et al., 2017; Tôrres & Diniz-Filho, 2004) with Spearman's correlation tests (Spearman, 1904). Using the 'picante' package (Kembel et al., 2010), we calculated PD for each group as the sum of branch lengths connecting a given set of species to the root of a phylogenetic tree (Faith, 1992).

Because PD is often correlated with richness, decoupling these two metrics provides a powerful tool to assess community phylogenetic structure and, thus, offers valuable insights into macroecological and biogeographical processes (Webb et al., 2002). We did this by running linear regression analyses in order to model the relationship between phylogenetic diversity and richness and then assessed the residuals of these models (PD $\sim$ SR residuals). When higher or lower than zero, PD $\sim$ SR residuals may indicate PD higher or lower than expected based on richness, respectively (Brown et al., 2020; Colville et al., 2020; Forest et al., 2007).

Also, to get a more comprehensive perspective on the spatial distribution patterns of richness, PD and PD $\sim$ SR residuals, we combined data from all groups. Because the phylogenetic trees were scaled by total tree length, PD was comparable between results. Thus, we obtained total richness and PD by summing these metrics for all groups in each cell. We used these data to run a linear regression analysis and then assessed PD $\sim$ SR residuals for all groups combined.

#### S1.8 Beta diversity

In order to evaluate whether the plant communities in the *campos rupestres* are taxonomically and/or phylogenetically structured, we conducted UPGMA analyses for each group using dissimilarity matrices based on Jaccard distance (beta diversity; Jaccard 1901) and UniFrac (phylobeta diversity; Lozupone & Knight 2005). As with alpha diversity metrics, we also performed an UPGMA analysis based on Jaccard distance for all groups combined simply by merging our distribution matrices. This approach was not possible with UniFrac, however, since this metric relies on pairwise calculations based on phylogenetic trees.

The consensus dendrograms were each based on 1000 replicates, with a consensus criterion of 0.5. Defining a cutting level in a dendrogram in order to visualise meaningful groups is highly arbitrary, but we based our decision on fusion level values (Borcard et al., 2018) and attempted to maintain a consistent number of clusters between all groups. Prior to conducting these analyses, we removed singletons (species whose occurrence is restricted to a single cell) in order to avoid data noise. Jaccard distance assesses the degree in which a given pair of communities are dissimilar to each other. Likewise, UniFrac gives the proportion of the phylogenetic tree which is not shared between two sites. These metrics provide values ranging from zero to one, with zero indicating complete similarity and one implying total dissimilarity. These analyses were conducted using the 'recluster' package (Dapporto et al., 2013).

We decided to use Jaccard distance and UniFrac to assess both present and past aspects of community assembling processes. While beta diversity provides useful information on the amount of taxonomic overlapping between sites, phylobeta diversity adds the dimension of evolutionary time into this context (Emerson & Gillespie, 2008; Graham & Fine, 2008; Wiens & Donoghue, 2004), providing useful knowledge on potential historical constraints.

#### S1.9 CANAPE

The categorical analyses of neo- and paleo-endemism were conducted following the methods developed by Mishler et al. (2014), which involve prior calculation of PE (Rosauer et al., 2009) and RPE (Mishler et al., 2014). CANAPE consists of a two-step statistical process that identifies areas of statistically significant neo- or paleo-endemism. We based this process on a null model which randomises occurrence matrices while fixing richness and frequency of occurrence (999 replicates for each instance). PE and RPE were calculated with the 'PDcalc'package by D. Nipperess (under development). We ran CANAPE according to the algorithm hereinafter specified.

First, to evaluate if a given grid cell harbours statistically significant phylogenetic endemism, either actual or comparative PE (i.e. PE based on a tree with equal branch lengths) must be significantly high (one-tailed test, $\alpha = 0,05$). If this condition holds, then the grid cell is subsequently defined as one of the non-overlapping categories of endemism distinguished by CANAPE. If RPE is significantly low or high (two-tailed test, $\alpha = 0,05$), the grid is respectively defined as a neo-endemic or paleo-endemic site. Otherwise, if this condition does not hold but both actual and comparative PE are significantly high (one-tailed test, $\alpha = 0,05$), the grid is a centre of mixed endemism, a category which has neither a prevalence of long, rare branches nor of short, rare ones, but a mix of both. A grid which is defined as such can be further categorised as a super endemic site if both actual and comparative PE are significantly high at a lower confidence interval ($\alpha = 0,01$). In this study, we found some cells for which the first condition held true (i.e. either actual or comparative PE were significantly high) but then did not fall in any of the categories CANAPE is able do distinguish. We defined those cells as harbouring an 'uncertain' endemism type.

Because we did not consider occurrence data within adjacent areas, we recognise that a potential bias in our analyses is that they are only relative to the *campos rupestres*. While this could be problematic regarding analyses of endemism, CANAPE relies on the range of phylogenetic branches rather than that of terminals. Besides, the *campos rupestres* are known to harbour a exceedingly high proportion of endemics (Giulietti et al., 1997), most of which are restricted to small areas (e.g. Bitencourt & Rapini, 2013; Echternacht et al., 2011; Rapini, 2010; Trovó et al., 2013). Thus, we believe our data and analyses were robust enough to assess the distribution of different types of endemism within the *campos rupestres*' range. Nevertheless, including the surrounding vegetation in future studies will certainly improve knowledge on these patterns.

### Appendix S2: Additional figures

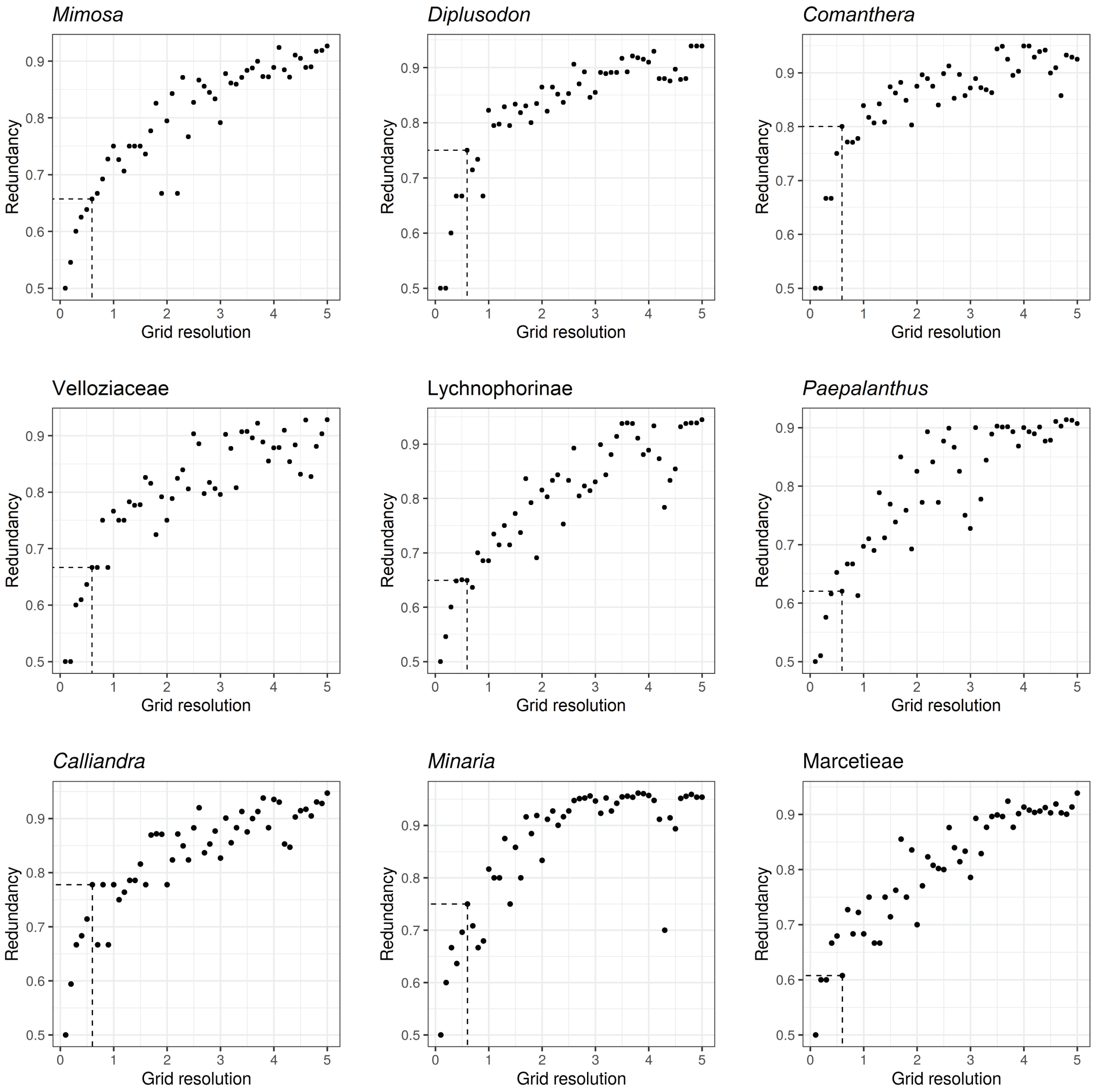


**Figure S2. 1:** Median values of redundancy based on data of the respective focal group plotted against grid resolution in decimal degrees. The higher redundancy is, the better the sampling quality. Note that redundancy for all groups steadily increases alongside grid cells' size, reaching a plateau at around 0.9. Dashed lines indicate redundancy for the chosen grid resolution (0.6 per 0.6 decimal degrees).

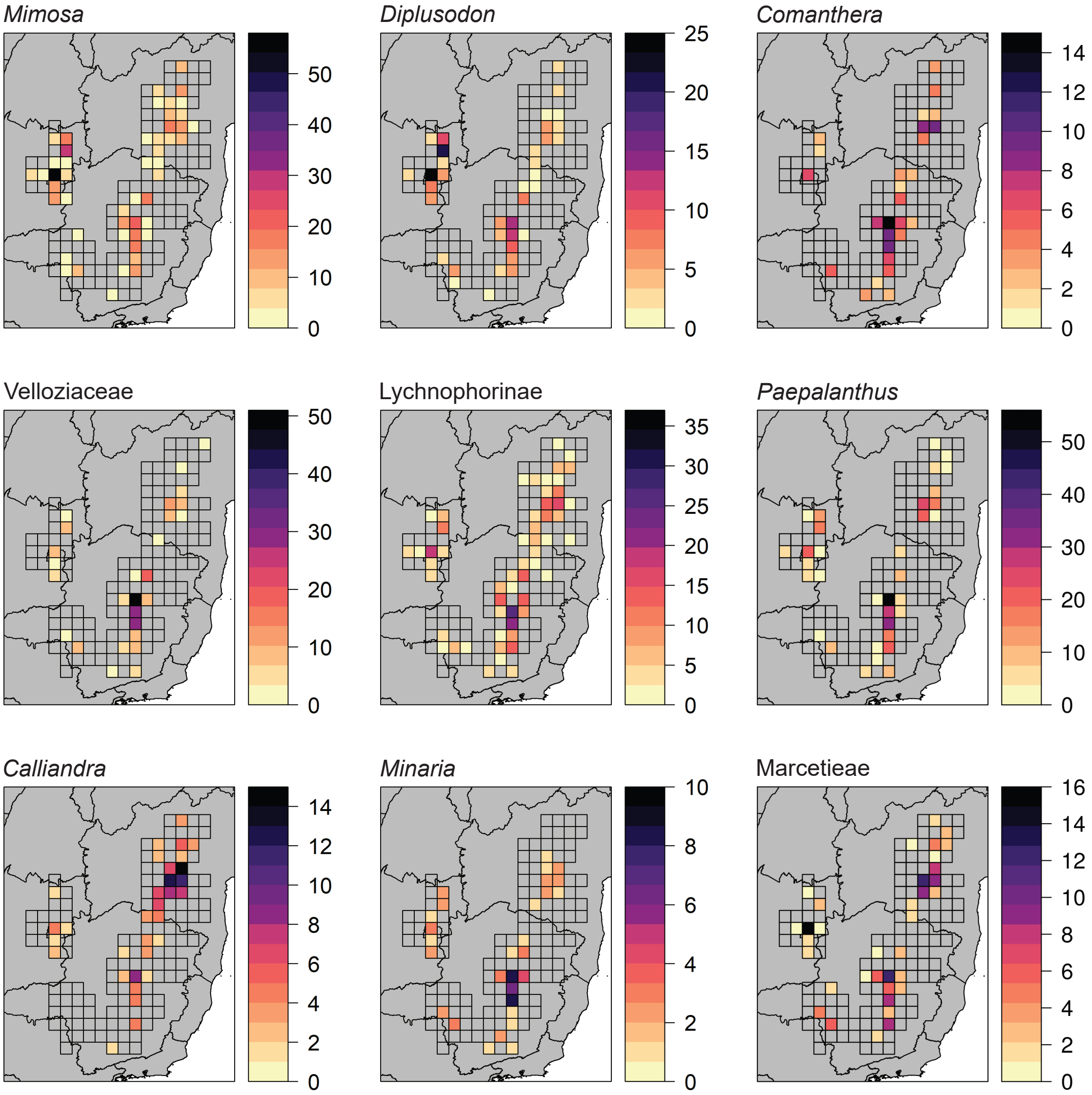


**Figure S2. 2:** Spatial distribution of species richness of each focal group across the campos rupestres. Transparent grid cells indicate absent data.

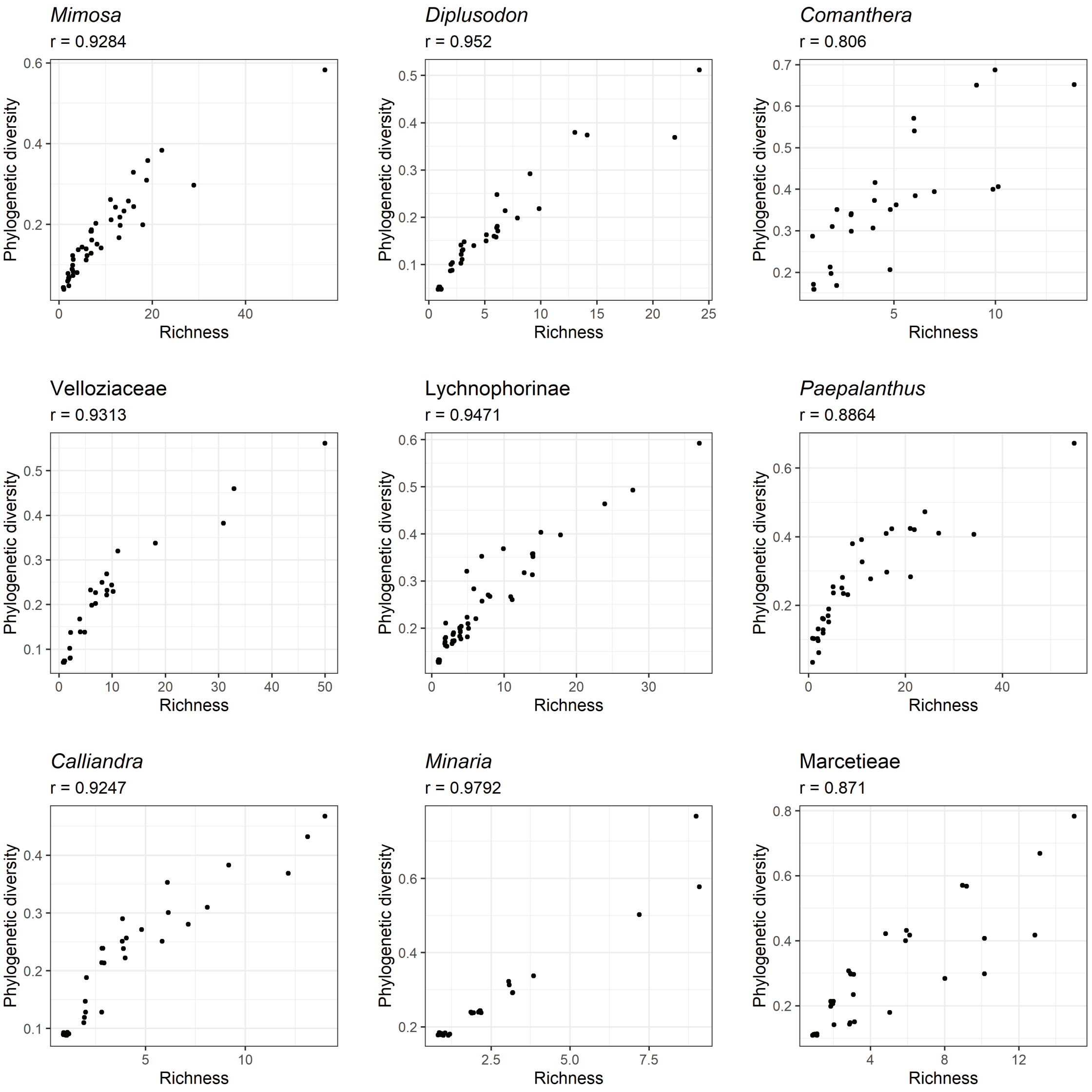


**Figure S2. 3:** Relationship between phylogenetic diversity and species richness based on data of the respective focal group. Spearman's correlation coefficients (r) indicate a strong and positive correlation.

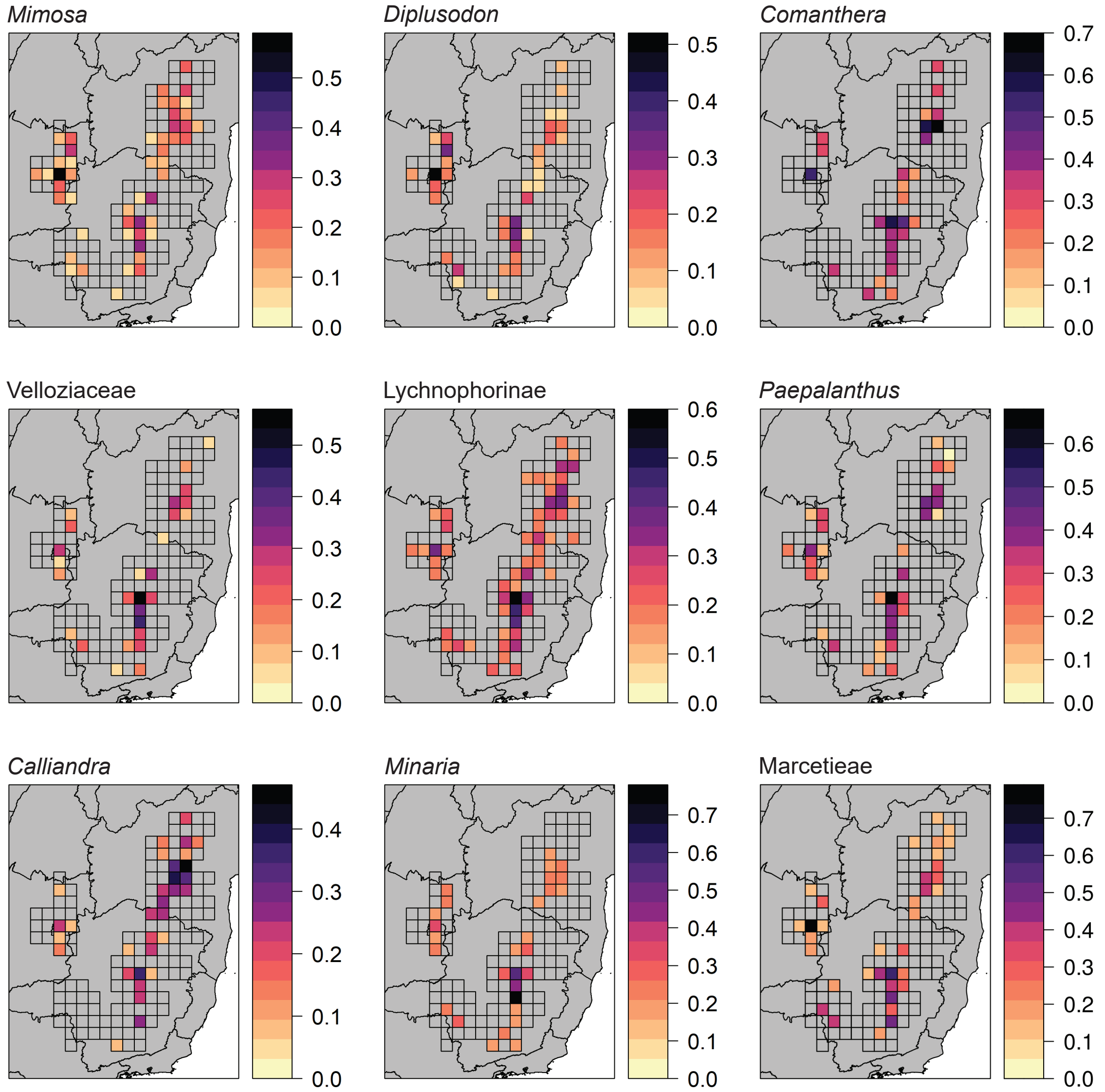


**Figure S2. 4:** Spatial distribution of phylogenetic diversity of each focal group across the campos rupestres. Transparent grid cells indicate absent data.

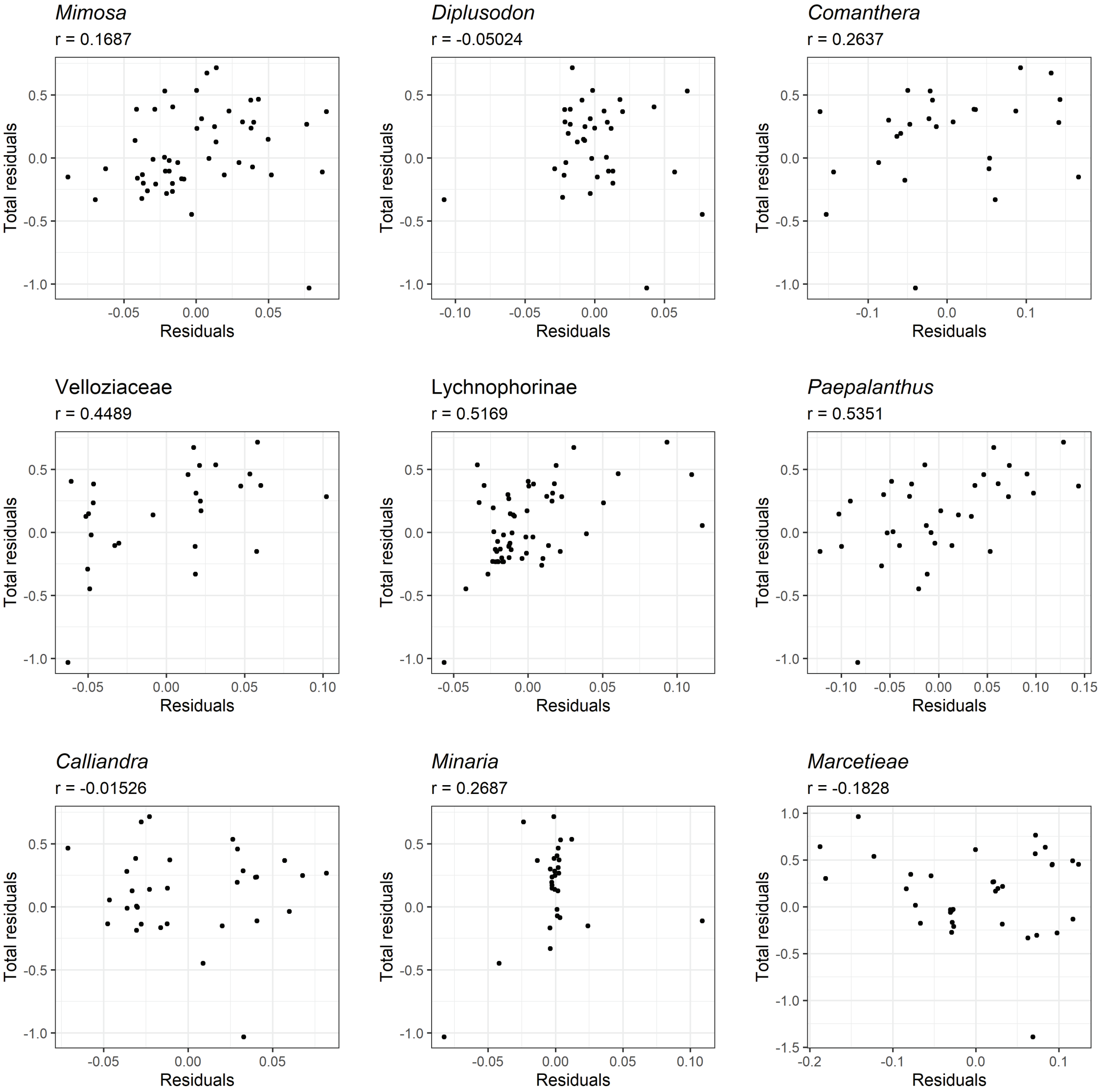


**Figure S2. 5:** Relationship between residuals of linear regressions using data from all groups combined (Total residuals) and data from the respective group (Residuals). Note that some groups (Velloziaceae, Lychnophorinae and Paepalanthus) contribute more than others to the distribution pattern of total residuals.

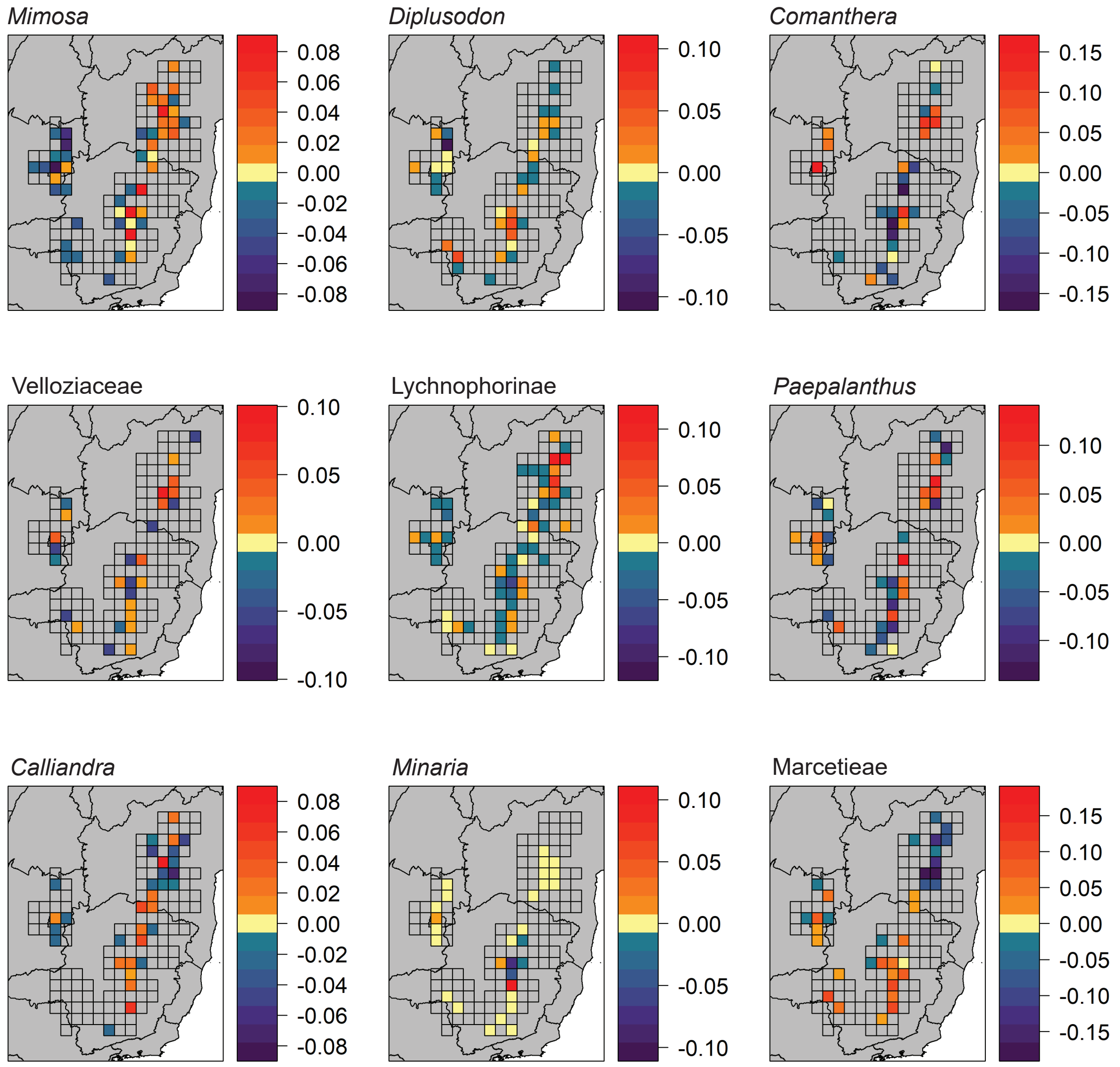


**Figure S2. 6:** Spatial distribution of residuals from linear regressions between species richness and phylogenetic diversity for each focal group across the campos rupestres. Positive and negative residuals respectively indicate phylogenetic overdispersion and clustering (i.e. communities in which phylogenetic diversity is higher or lower than expected based on species richness). Transparent grid cells indicate absent data.

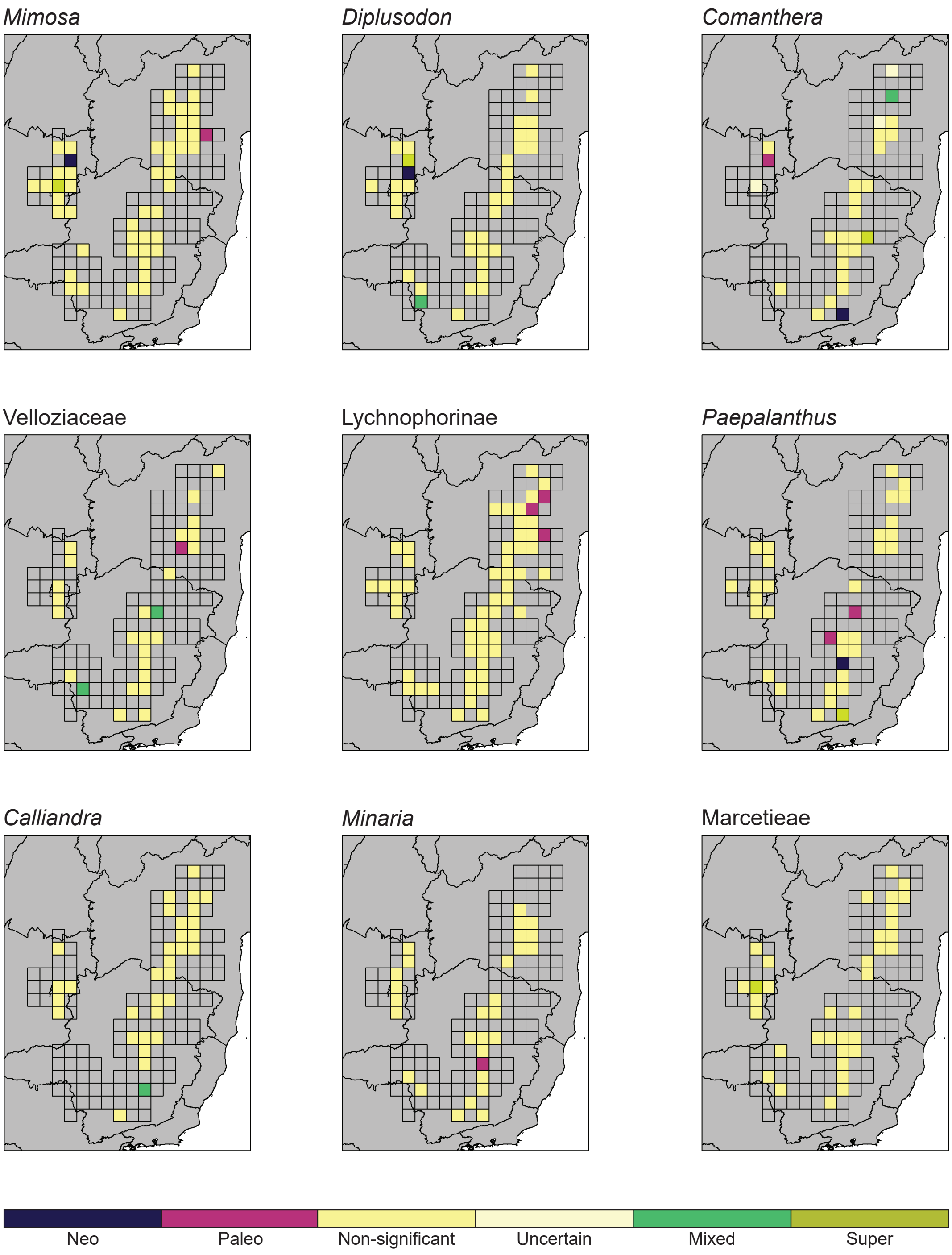


**Figure S2. 7:** CANAPE results. Neo-endemic cells contain a predominance of short, rare phylogenetic branches; paleo-endemic cells contain a predominance of long, rare branches; and cells with mixed endemism contain a mixture of both. Mixed endemism can be further categorized as super endemism in a cell in which it is statistically significant at a lower confidence interval ($\alpha \leq 0.001$). Some cells in Comanthera passed the first statistical test that defines CANAPE, but then did not fall in any endemic category this analysis is able to distinguish. We defined those cells as harbouring an "uncertain" endemism type. Transparent grid cells indicate absent data.

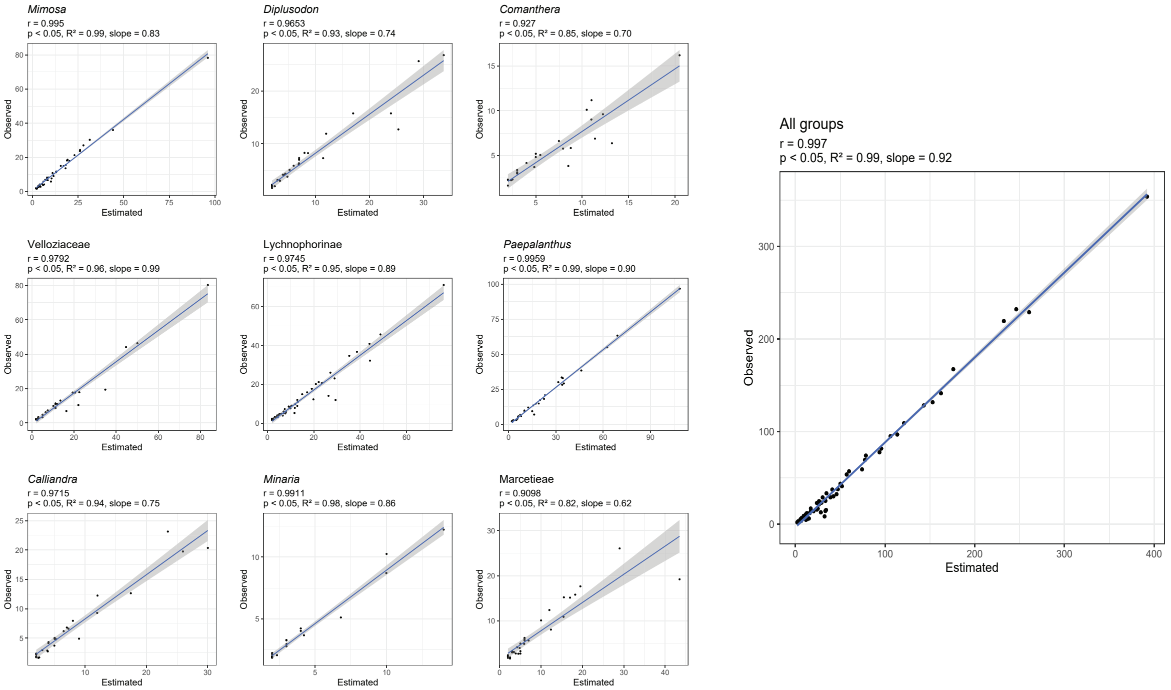


**Figure S2. 8:** Relationship between observed species richness (Observed) and asymptotic speciess richness (Estimated) estimated using a protocol based on Hill numbers. Pearson correlation (r) and model adjustment level ($R^2$) are high for all groups individually and combined. P-values show statistical significance for this relationship. Regression slopes lower than 1 indicate that estimated species richness is lower than observed species richness.

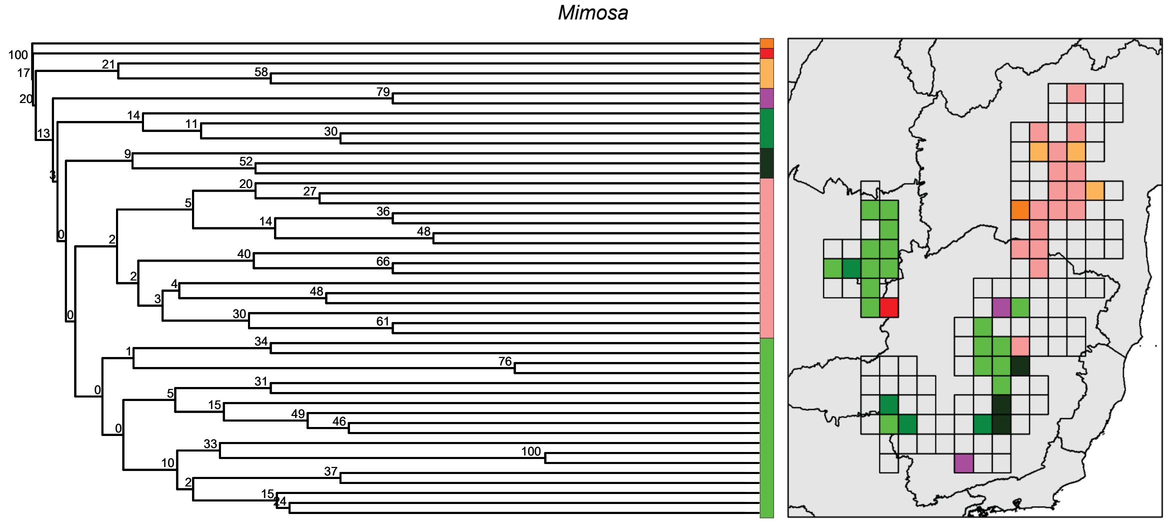


**Figure S2. 9:** Clustering support values (bootstrap) for the beta diversity (Jaccard) analysis in *Mimosa*. Bootstrap values were retrieved by resampling species occurring in each site (1000 iterations). Note that bootstrap values tend to decrease when using matrices of unequal distribution, which is generally the case of occurrence data. Therefore, low bootstrap values are often retrieved and are not very informative.

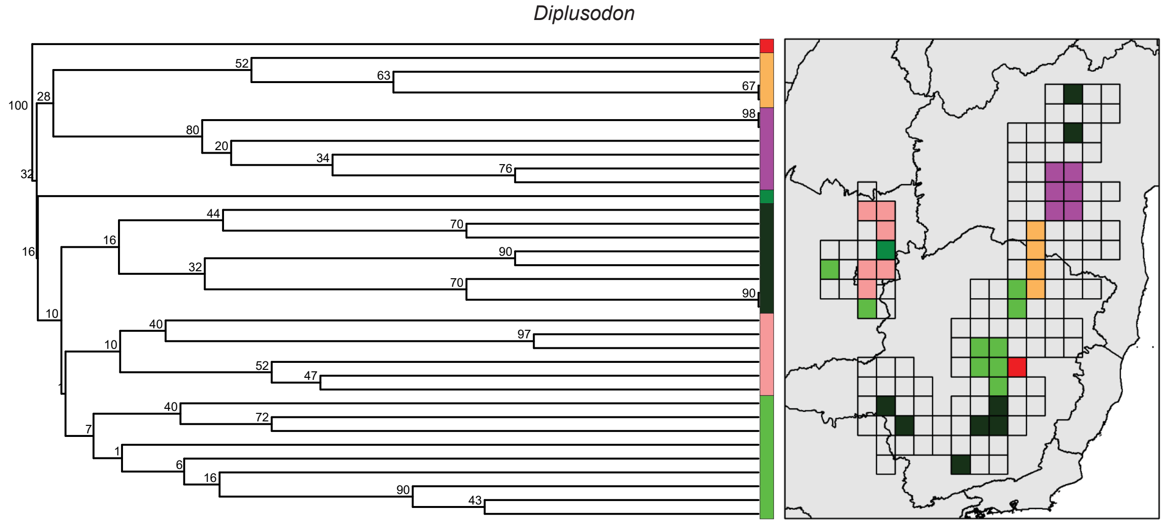


**Figure S2. 10:** Clustering support values (bootstrap) for the beta diversity (Jaccard) analysis in *Diplusodon*. Bootstrap values were retrieved by resampling species occurring in each site (1000 iterations). Note that bootstrap values tend to decrease when using matrices of unequal distribution, which is generally the case of occurrence data. Therefore, low bootstrap values are often retrieved and are not very informative.

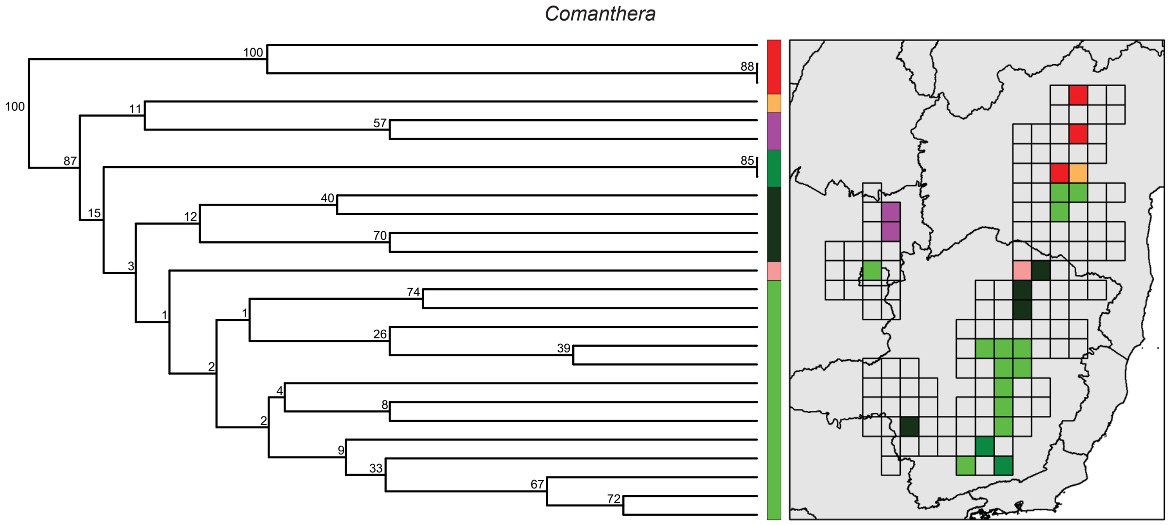


**Figure S2. 11:** Clustering support values (bootstrap) for the beta diversity (Jaccard) analysis in *Comanthera*. Bootstrap values were retrieved by resampling species occurring in each site (1000 iterations). Note that bootstrap values tend to decrease when using matrices of unequal distribution, which is generally the case of occurrence data. Therefore, low bootstrap values are often retrieved and are not very informative.

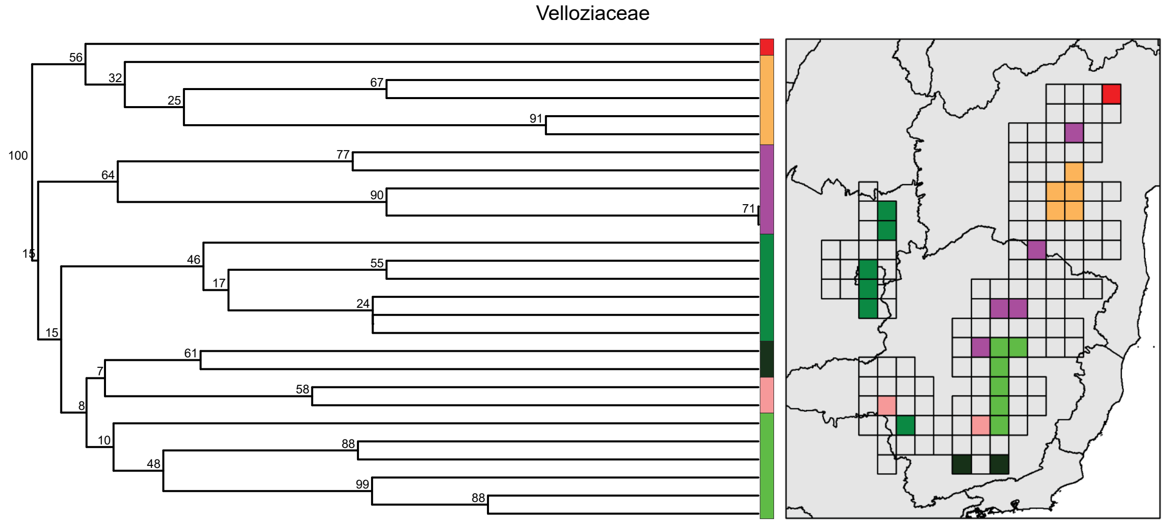


**Figure S2. 12:** Clustering support values (bootstrap) for the beta diversity (Jaccard) analysis in Velloziaceae. Bootstrap values were retrieved by resampling species occurring in each site (1000 iterations). Note that bootstrap values tend to decrease when using matrices of unequal distribution, which is generally the case of occurrence data. Therefore, low bootstrap values are often retrieved and are not very informative.

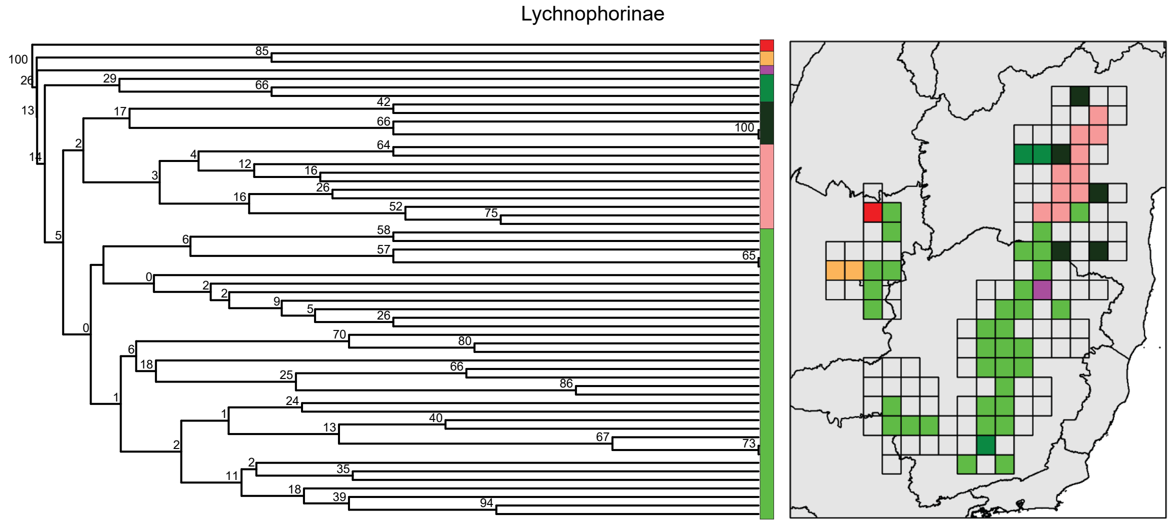


**Figure S2. 13:** Clustering support values (bootstrap) for the beta diversity (Jaccard) analysis in Lychnophorinae. Bootstrap values were retrieved by resampling species occurring in each site (1000 iterations). Note that bootstrap values tend to decrease when using matrices of unequal distribution, which is generally the case of occurrence data. Therefore, low bootstrap values are often retrieved and are not very informative.

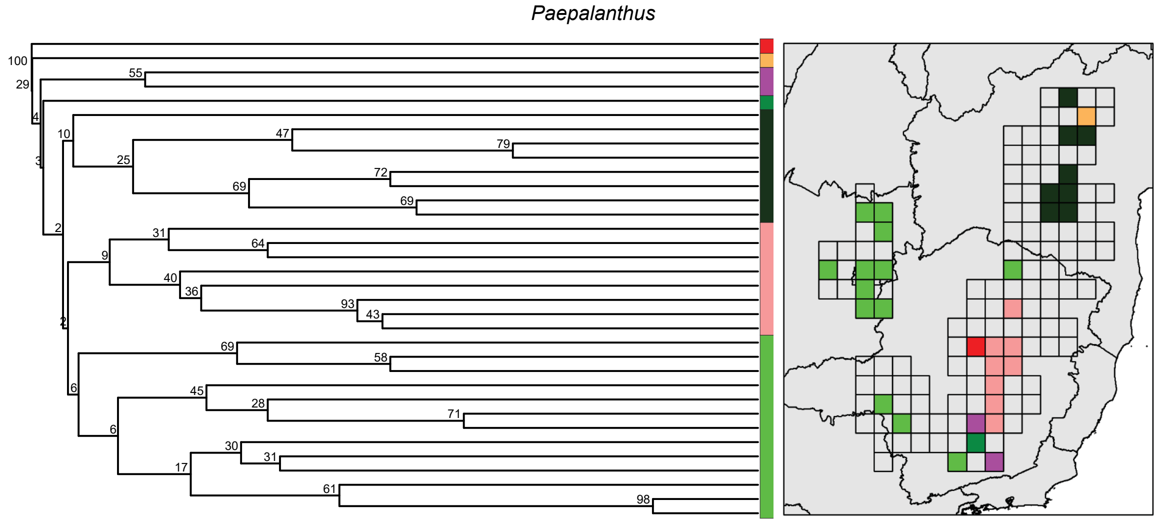


**Figure S2. 14:** Clustering support values (bootstrap) for the beta diversity (Jaccard) analysis in *Paepalanthus*. Bootstrap values were retrieved by resampling species occurring in each site (1000 iterations). Note that bootstrap values tend to decrease when using matrices of unequal distribution, which is generally the case of occurrence data. Therefore, low bootstrap values are often retrieved and are not very informative.

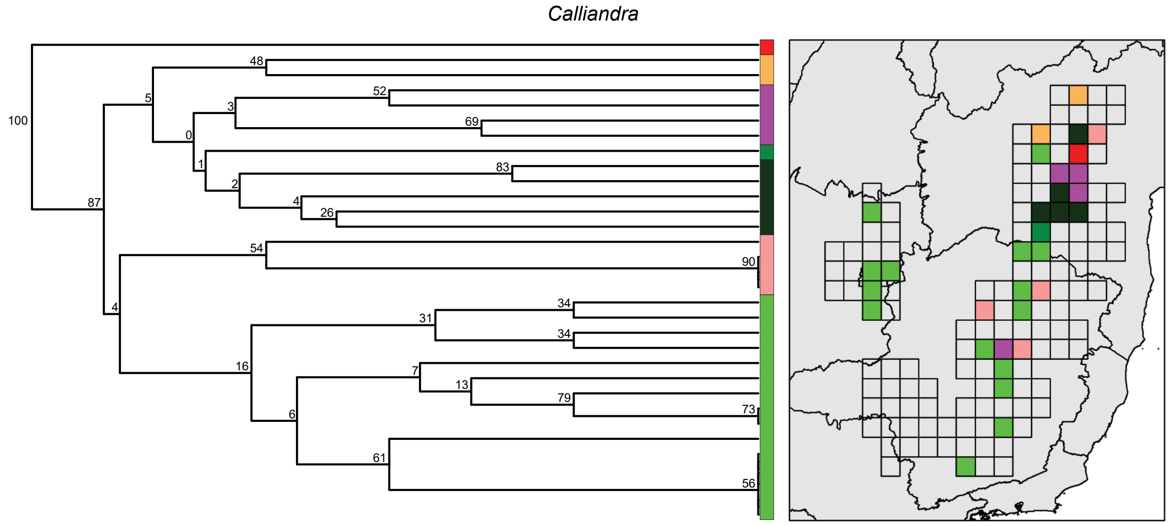


**Figure S2. 15:** Clustering support values (bootstrap) for the beta diversity (Jaccard) analysis in *Calliandra*. Bootstrap values were retrieved by resampling species occurring in each site (1000 iterations). Note that bootstrap values tend to decrease when using matrices of unequal distribution, which is generally the case of occurrence data. Therefore, low bootstrap values are often retrieved and are not very informative.

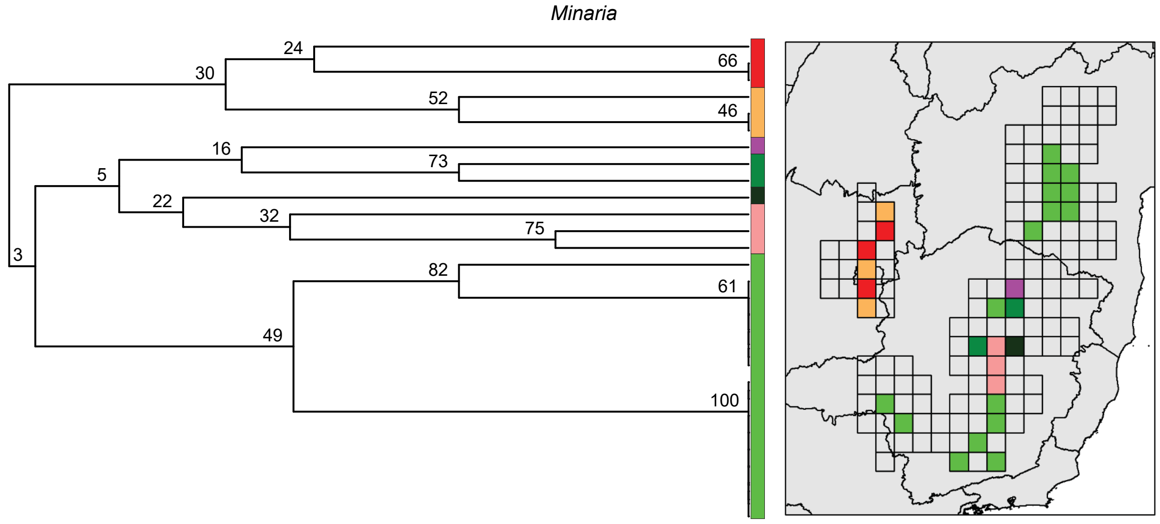


**Figure S2. 16:** Clustering support values (bootstrap) for the beta diversity (Jaccard) analysis in *Minaria*. Bootstrap values were retrieved by resampling species occurring in each site (1000 iterations). Note that bootstrap values tend to decrease when using matrices of unequal distribution, which is generally the case of occurrence data. Therefore, low bootstrap values are often retrieved and are not very informative.

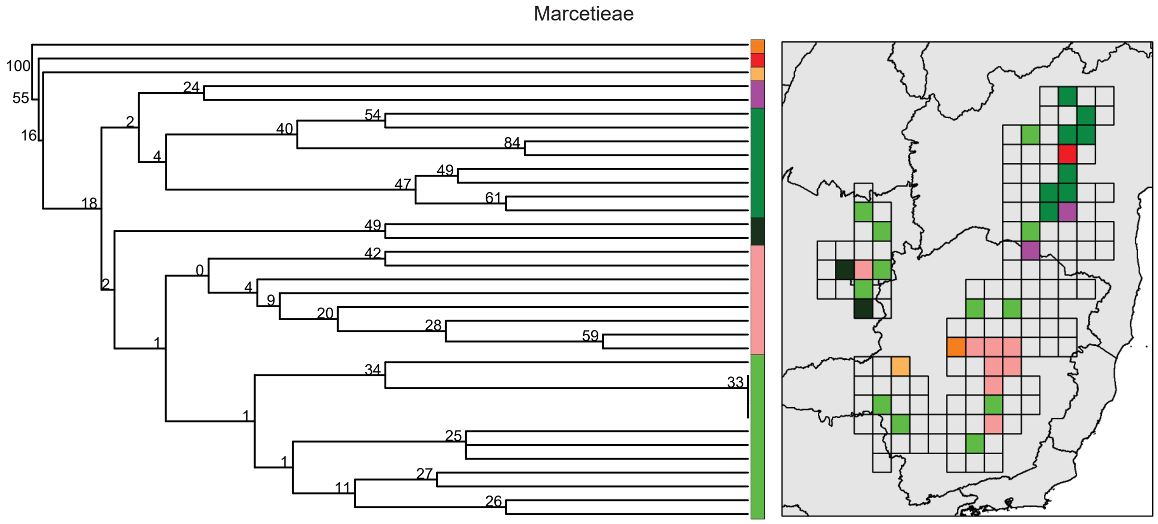


**Figure S2. 17:** Clustering support values (bootstrap) for the beta diversity (Jaccard) analysis in Marcetieae. Bootstrap values were retrieved by resampling species occurring in each site (1000 iterations). Note that bootstrap values tend to decrease when using matrices of unequal distribution, which is generally the case of occurrence data. Therefore, low bootstrap values are often retrieved and are not very informative.

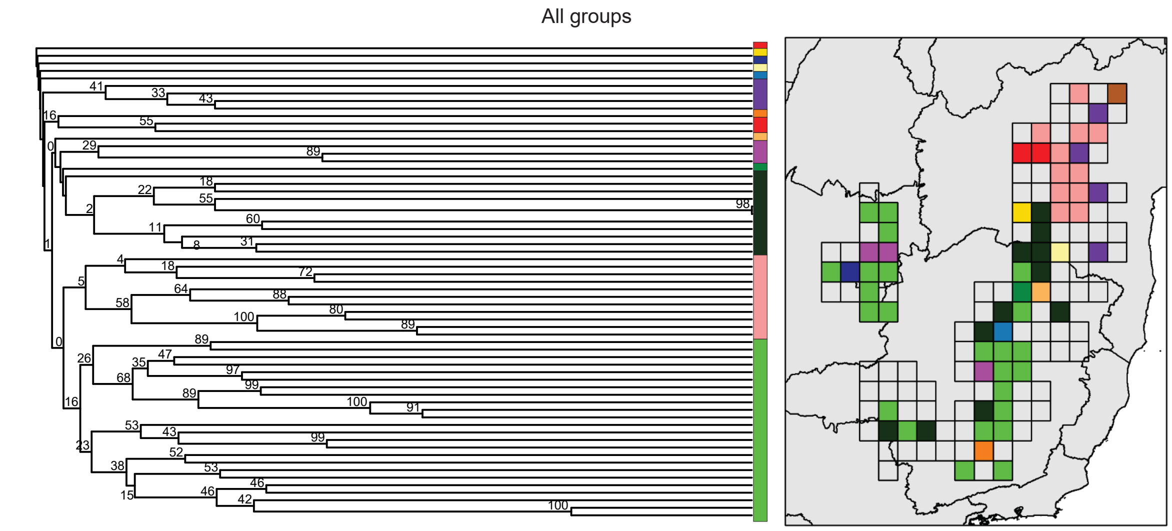


**Figure S2. 18:** Clustering support values (bootstrap) for the beta diversity (Jaccard) analysis in All groups. Bootstrap values were retrieved by resampling species occurring in each site (1000 iterations). Note that bootstrap values tend to decrease when using matrices of unequal distribution, which is generally the case of occurrence data. Therefore, low bootstrap values are often retrieved and are not very informative.

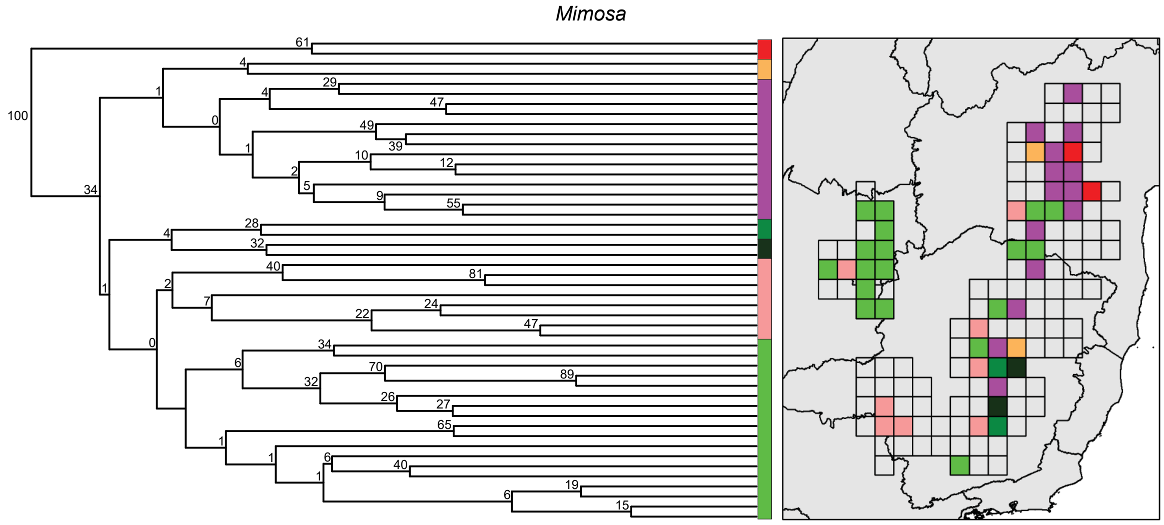


**Figure S2. 19:** Clustering support values (bootstrap) for the phylobeta diversity (UniFrac) analysis in *Mimosa*. Bootstrap values were retrieved by resampling species occurring in each site (1000 iterations). Note that bootstrap values tend to decrease when using matrices of unequal distribution, which is generally the case of occurrence data. Therefore, low bootstrap values are often retrieved and are not very informative.

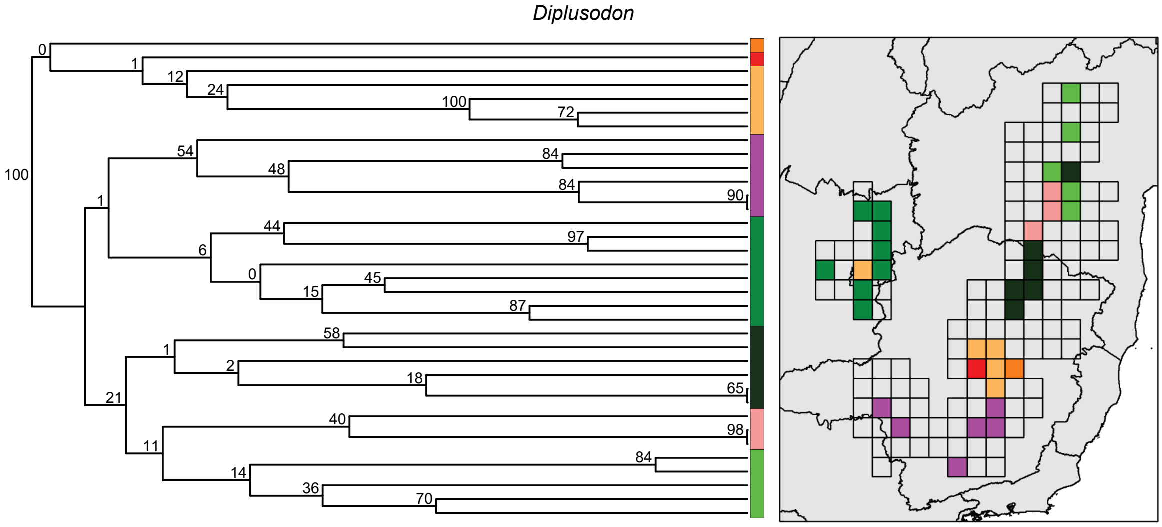


**Figure S2. 20:** Clustering support values (bootstrap) for the phylobeta diversity (UniFrac) analysis in *Diplusodon*. Bootstrap values were retrieved by resampling species occurring in each site (1000 iterations). Note that bootstrap values tend to decrease when using matrices of unequal distribution, which is generally the case of occurrence data. Therefore, low bootstrap values are often retrieved and are not very informative.

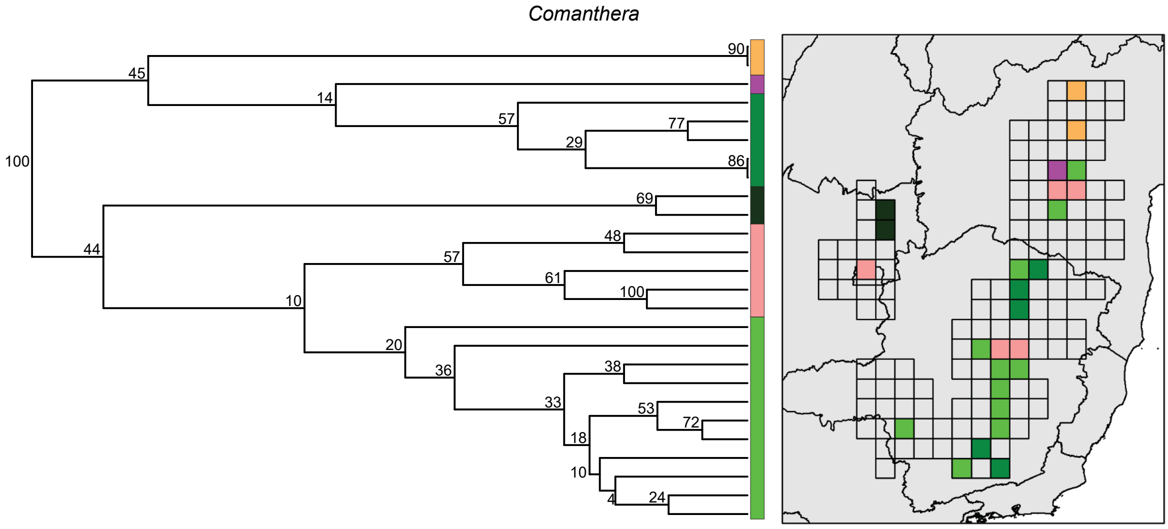


**Figure S2. 21:** Clustering support values (bootstrap) for the phylobeta diversity (UniFrac) analysis in *Comanthera*. Bootstrap values were retrieved by resampling species occurring in each site (1000 iterations). Note that bootstrap values tend to decrease when using matrices of unequal distribution, which is generally the case of occurrence data. Therefore, low bootstrap values are often retrieved and are not very informative.

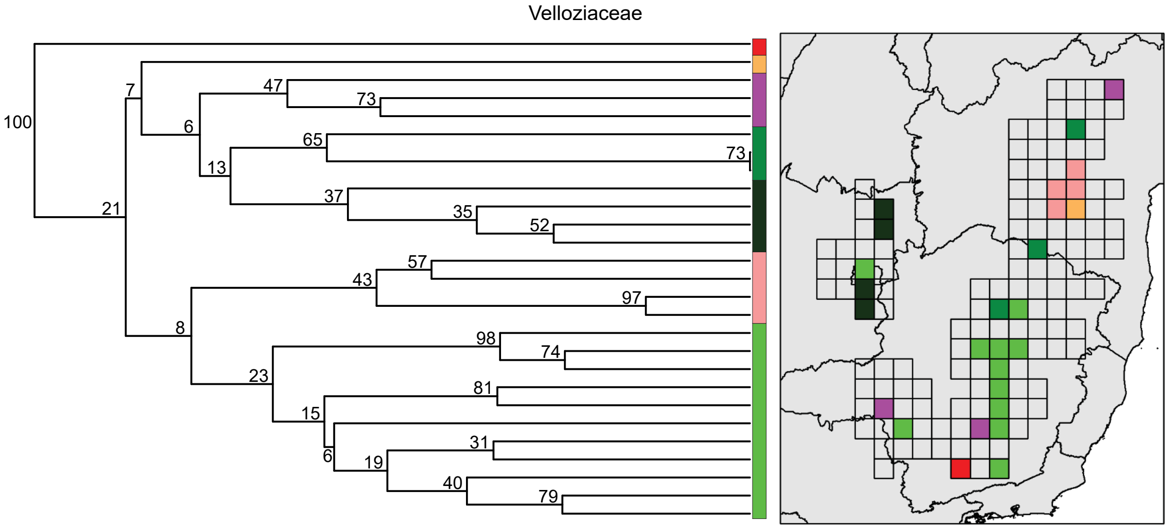


**Figure S2. 22:** Clustering support values (bootstrap) for the phylobeta diversity (UniFrac) analysis in Velloziaceae. Bootstrap values were retrieved by resampling species occurring in each site (1000 iterations). Note that bootstrap values tend to decrease when using matrices of unequal distribution, which is generally the case of occurrence data. Therefore, low bootstrap values are often retrieved and are not very informative.

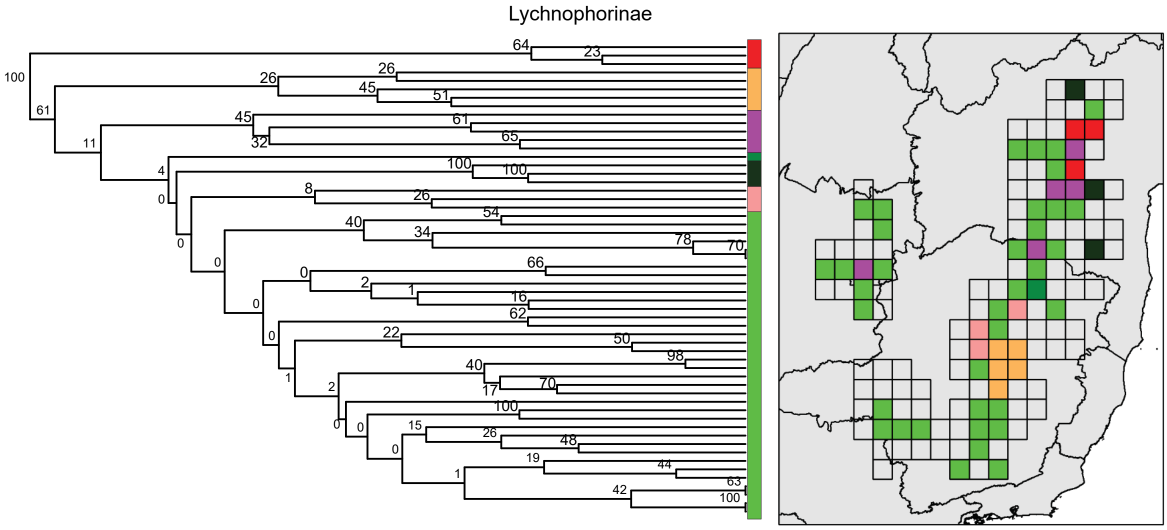


**Figure S2. 23:** Clustering support values (bootstrap) for the phylobeta diversity (UniFrac) analysis in Lychnophorinae. Bootstrap values were retrieved by resampling species occurring in each site (1000 iterations). Note that bootstrap values tend to decrease when using matrices of unequal distribution, which is generally the case of occurrence data. Therefore, low bootstrap values are often retrieved and are not very informative.

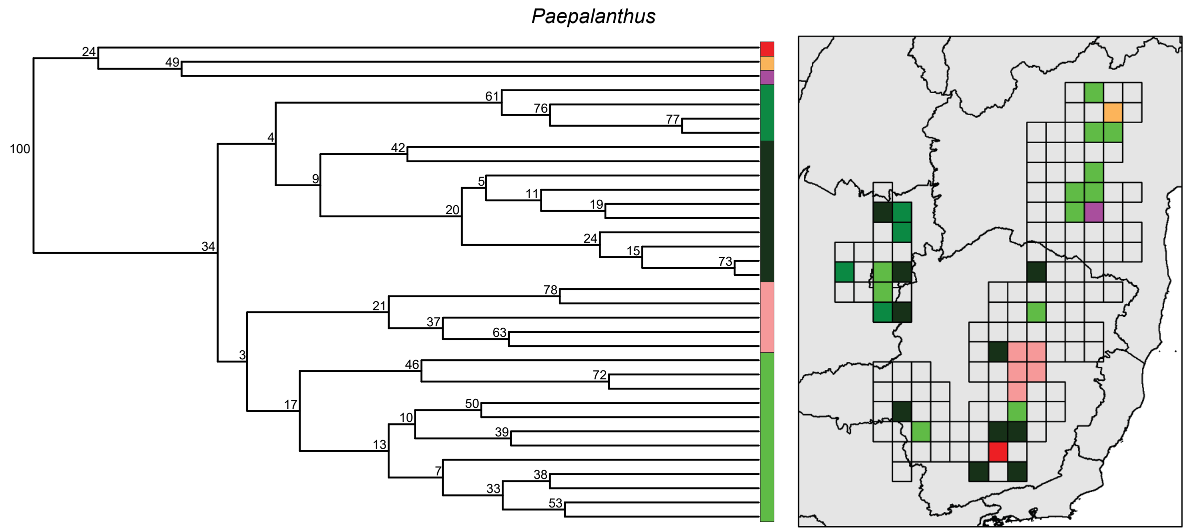


**Figure S2. 24:** Clustering support values (bootstrap) for the phylobeta diversity (UniFrac) analysis in *Paepalanthus*. Bootstrap values were retrieved by resampling species occurring in each site (1000 iterations). Note that bootstrap values tend to decrease when using matrices of unequal distribution, which is generally the case of occurrence data. Therefore, low bootstrap values are often retrieved and are not very informative.

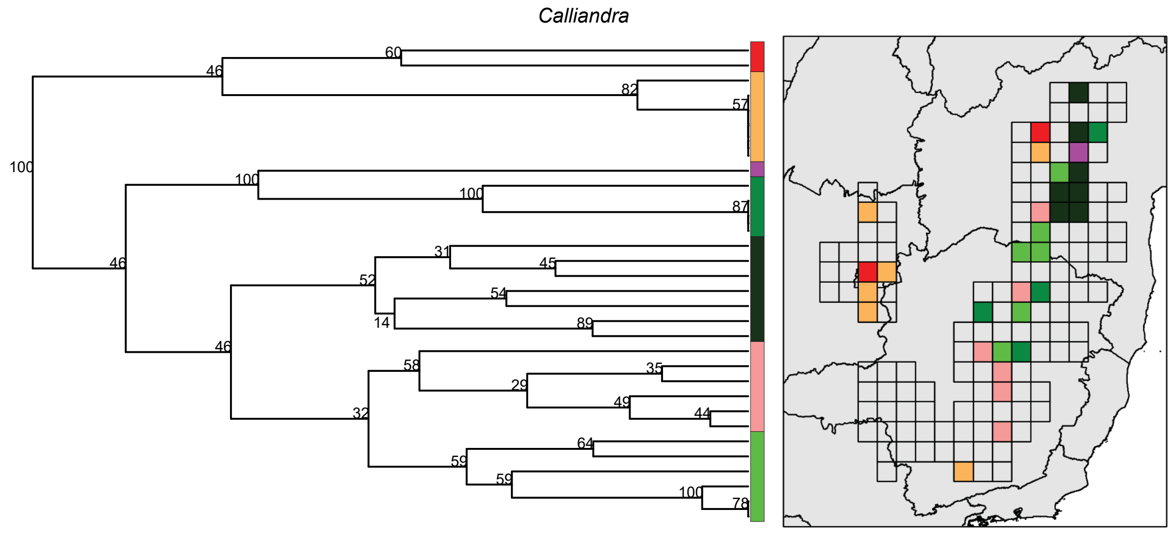


**Figure S2. 25:** Clustering support values (bootstrap) for the phylobeta diversity (UniFrac) analysis in *Calliandra*. Bootstrap values were retrieved by resampling species occurring in each site (1000 iterations). Note that bootstrap values tend to decrease when using matrices of unequal distribution, which is generally the case of occurrence data. Therefore, low bootstrap values are often retrieved and are not very informative.

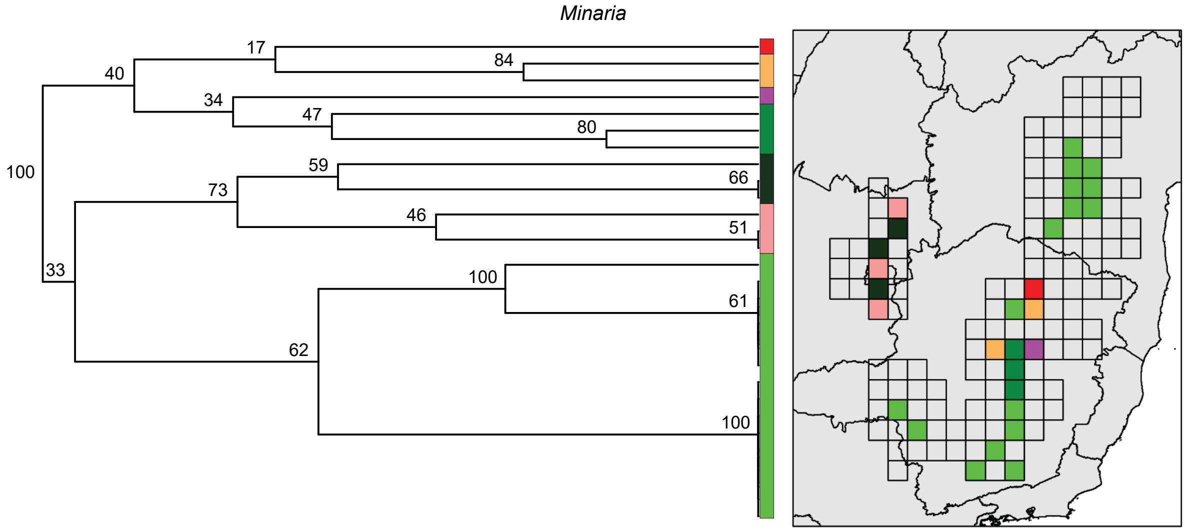


**Figure S2. 26:** Clustering support values (bootstrap) for the phylobeta diversity (UniFrac) analysis in *Minaria*. Bootstrap values were retrieved by resampling species occurring in each site (1000 iterations). Note that bootstrap values tend to decrease when using matrices of unequal distribution, which is generally the case of occurrence data. Therefore, low bootstrap values are often retrieved and are not very informative.

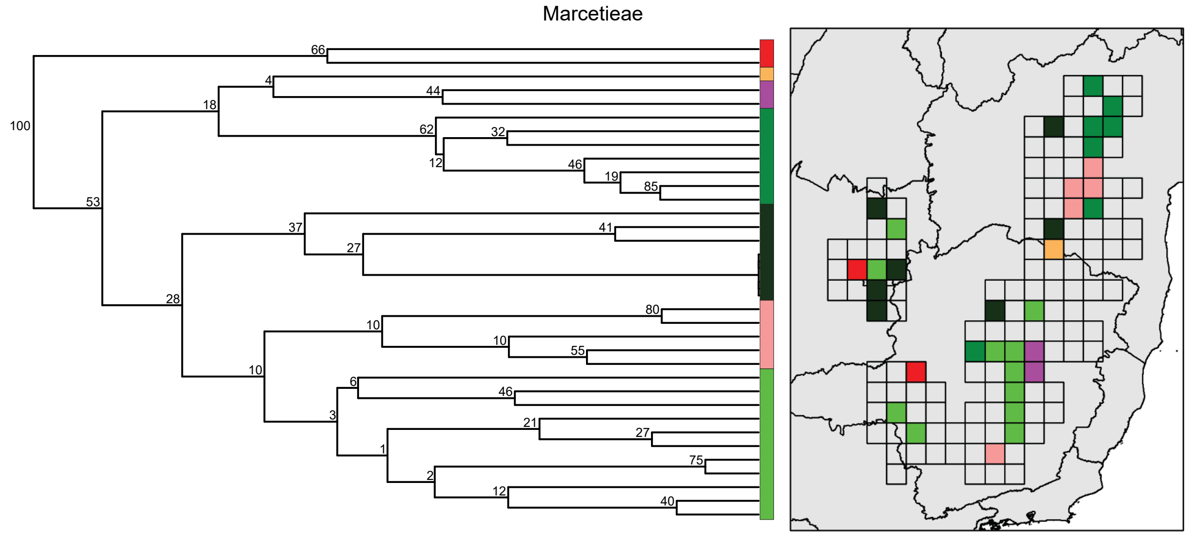


**Figure S2. 27:** Clustering support values (bootstrap) for the phylobeta diversity (UniFrac) analysis in Marcetieae. Bootstrap values were retrieved by resampling species occurring in each site (1000 iterations). Note that bootstrap values tend to decrease when using matrices of unequal distribution, which is generally the case of occurrence data. Therefore, low bootstrap values are often retrieved and are not very informative.
